# Interaction between induced and natural variation at *oil yellow1* delays reproductive maturity in maize

**DOI:** 10.1101/706846

**Authors:** Rajdeep S. Khangura, Bala P. Venkata, Sandeep R. Marla, Michael V. Mickelbart, Singha Dhungana, David M. Braun, Brian P. Dilkes, Gurmukh S. Johal

## Abstract

We previously demonstrated that maize (*Zea mays*) locus *very oil yellow1 (vey1)* encodes a putative cis-regulatory expression polymorphism at the magnesium chelatase subunit I gene (aka *oil yellow1*) that strongly modifies the chlorophyll content of the semi-dominant *Oy1-N1989* mutants. The *vey1* allele of Mo17 inbred line reduces chlorophyll content in the mutants leading to reduced photosynthetic output. *Oy1-N1989* mutants in B73 reached reproductive maturity four days later than wild-type siblings. Enhancement of *Oy1-N1989* by the Mo17 allele at the *vey1* QTL delayed maturity further, resulting in detection of a flowering time QTL in two bi-parental mapping populations crossed to *Oy1-N1989*. The near isogenic lines of B73 harboring the *vey1* allele from Mo17 delayed flowering of *Oy1-N1989* mutants by twelve days. Just as previously observed for chlorophyll content, *vey1* had no effect on reproductive maturity in the absence of the *Oy1-N1989* allele. Loss of chlorophyll biosynthesis in *Oy1-N1989* mutants and enhancement by *vey1* reduced CO_2_ assimilation. We attempted to separate the effects of photosynthesis on the induction of flowering from a possible impact of chlorophyll metabolites and retrograde signaling by manually reducing leaf area. Removal of leaves, independent of the *Oy1-N1989* mutant, delayed flowering but surprisingly reduced chlorophyll contents of emerging leaves. Thus, defoliation did not completely separate the identity of the signal(s) that regulates flowering time from changes in chlorophyll content in the foliage. These findings illustrate the necessity to explore the linkage between metabolism and the mechanisms that connect it to flowering time regulation.

## Introduction

The onset of flowering in angiosperms has been a key focus for plant biologists working on ornamental, horticultural, and other crop species (Lang 1952; Zeevaart 1962; Searle 1965). The onset of reproductive development in angiosperms is affected by a change in meristem identity. The vegetative to floral transition of meristems commits plant development to production of floral organs and sexual reproduction. The integration of signals to correctly time this transition is key to plant fitness. Unsurprisingly, endogenous and environmental cues regulate flowering time (Amasino and Michaels 2010; Cho *et al.* 2017; Minow *et al.* 2018). A critical environmental cue is the duration of the light period, or photoperiod. The photoperiodic responses of plants influence the vegetative to floral transition and the mechanisms of this response have been a focus of intensive research for over a century (Klebs, 1918). Multiple non-photoperiodic cues as well as endogenous signals, sometimes called the autonomous pathway, are also critical to floral transition. Endogenous signals, including hormones and the carbohydrate status of the plant, can also play a critical role in the regulation of flowering time (Corbesier *et al.* 1998; Moghaddam and Ende 2013). But it can be difficult to fully separate endogenous and environmental influences as some environmental factors, such as light quality, alter hormone biosynthesis (Lang 1957; Evans and Poethig 1995; Mutasa-Göttgens and Hedden 2009), and light powers photosynthesis and thereby carbohydrate status (Chen *et al.* 2004). These stimuli converge through the same floral integrators (named FLOWERING LOCUS T (FT) and FLOWERING LOCUS D (FD) in *Arabidopsis thaliana)* for which orthologs have been identified in many flowering plants (Abe *et al.* 2005; Wigge *et al.* 2005; Corbesier *et al.* 2007; Meng *et al.* 2011; Zhang *et al.* 2016). Accumulation of FT and FD gene products trigger the vegetative shoot apical meristems to acquire the competency to become inflorescence meristems and produce flowers in part via the activation of MADS-box transcription factors that control meristem identity through APETALA1 (Abe *et al.* 2005; Wigge *et al.* 2005). In Arabidopsis, FT is regulated by CONSTANS (CO) in response to both circadian regulation and photoperiodic responses, and CO regulates the MADS-box transcription factor SUPPRESSOR OF CONSTANS1 (SOC1) through FT (Samach *et al.* 2000; Yoo *et al.* 2005).

Maize was domesticated from teosinte (*Zea mays* ssp. *parviglumis* or spp. *mexicana*) in Central America (Doebley *et al.* 1997; Wang *et al.* 1999). Strong selection on time to reproductive maturity contributed to the adaptation of maize to different latitudes (Salvi *et al.* 2007; Huang *et al.* 2017; Swarts *et al.* 2017). Flowering of teosinte is promoted by short-day conditions. In contrast, temperate maize germplasm is relatively day-neutral and flowering is primarily under the control of the autonomous pathway (Coles *et al.* 2010). Mutant studies have identified loci critical to flowering in maize including: *indeterminate1* (*id1*; Colasanti et al. 1998); *early phase change* (*epc*; Vega et al. 2002); *delayed flowering1* (*dlf1*; Muszynski et al. 2006); the cis-element polymorphism *vegetative transitioning1* (*vgt1*; Salvi et al. 2007) that regulates a downstream APETALA2-like transcription factor *zmrap2.7*; *zea mays mads4* (*zmm4*; Danilevskaya et al. 2008); *zmcct10* (Hung *et al.* 2012); *zea mays centroradiales8* (*zcn8*; Meng et al. 2011); *zea mays mads1* (*zmmads1*; Alter et al. 2016); and *zea mays mads69* (Liang *et al.* 2019). Many of these loci encode the maize orthologs of genes identified as regulators of flowering in Arabidopsis. For example, *dlf1* and *zcn8* encode homologs of the Arabidopsis flowering time determinants FD and FT, respectively. *id1* encodes a zinc-finger transcription factor acting upstream of both DLF1 and ZCN8 (Kozaki *et al.* 2004; Muszynski *et al.* 2006; Meng *et al.* 2011). *zmm4* is an activator of flowering that is part of a conserved syntenic pair of MADS box genes in the grasses, with *zmm24* as the neighboring gene, and encodes one of two maize paralogs of the wheat flowering time and vernalization response locus VRN1 (Danilevskaya *et al.* 2008). *zmm4* acts downstream of *dlf1* and *id1* in the control of flowering time in maize. *zmmads1* is a functional homolog of the Arabidopsis flowering time and circadian rhythm regulator *soc1* (Alter *et al.* 2016). Several QTL studies have used the convenient phenotype of days to reproductive maturity as a proxy for flowering time and identified alleles controlling this trait in maize (Buckler *et al.* 2009; Coles *et al.* 2010; Steinhoff *et al.* 2012; Bouchet *et al.* 2017). While this trait is convenient it is determined by both the days to floral transition of the meristem and the growth rate of the stem and emergence and maturation of floral structures. Nevertheless, many natural variants controlling days to reproductive maturity in maize map to *bona fide* flowering time loci identified by mutant studies including alleles of *zmmads69*, *zmcct10*, *zcn8*, *dlf1*, and *vgt1* (Muszynski *et al.* 2006; Salvi *et al.* 2007; Meng *et al.* 2011; Hung *et al.* 2012; Guo *et al.* 2018; Liang *et al.* 2019).

One important endogenous signal that contributes to flowering time is the carbohydrate allocation status (Ohto *et al.* 2001; Seo *et al.* 2011; Eveland and Jackson 2012; Wahl *et al.* 2013; Cho *et al.* 2018). In maize, mutants that are compromised in either sugar export from source tissues or loading sucrose into the phloem flower later than their congenic wild-type siblings (Braun *et al.* 2006; Baker and Braun 2008; Ma *et al.* 2008; Slewinski *et al.* 2009; Slewinski and Braun 2010). This is not limited to maize, as starch-deficient Arabidopsis mutants exhibit delayed flowering (Corbesier *et al.* 1998). Trehalose-6-phosphate (T6P) has been implicated as a reporter of the energy status and Arabidopsis mutants effected in this metabolite also exhibit altered flowering time (Paul 2008; Wahl et al. 2013; Seo et al. 2011). T6P and sucrose are positively correlated, and low levels of T6P results in delayed flowering in Arabidopsis (Wahl *et al.* 2013). Remarkably, the carbohydrate status and transcriptional regulatory genes controlling flowering time may be directly linked in maize. The *id1* flowering time mutants alter carbohydrate partitioning in leaves and accumulate more sucrose and starch (Coneva et al. 2012). As a result, ID1 has been proposed to act as a carbohydrate status sensor that influences flowering time in maize (Coneva, Zhu, and Colasanti 2007; Minow et al. 2018). Remarkably, the promoter of the T6P biosynthetic gene *trehalose 6-phosphate synthase1* (*tps1*) is a predicted target of the ID1 DNA-binding protein and low levels of T6P were observed in *id1* mutants (Minow *et al.* 2018).

If sugars are critical for floral transitioning in plants, then manipulation of photosynthesis should alter flowering. Magnesium chelatase (MgChl) is a hetero-oligomeric enzyme complex comprised of subunits I, D, and H. This enzyme catalyzes the first committed step of chlorophyll biosynthesis by conversion of protoporphyrin IX (PPIX) into magnesium-PPIX (Walker and Weinstein 1991; Gibson et al. 1995). The I subunit of MgChl is encoded by *oil yellow1* (*oy1*) in maize (Sawers *et al.* 2006). The OY1-N1989 mutant protein carries a L176F amino acid substitution that results in the protein acting as a competitive inhibitor of MgChl complex function, and decouples ATPase and Mg^2+^ chelatase activity (Hansson *et al.* 1999, 2002; Sawers *et al.* 2006). As a result, homozygous *Oy1-N1989* mutants are seedling lethal with no chlorophyll accumulation but are viable to reproductive maturity in heterozygous condition (Sawers *et al.* 2006, Khangura *et al.* 2019).

We previously identified a cis-acting expression polymorphism at the *oy1* locus associated with a QTL called *very oil yellow1* (*vey1*) that modifies the chlorophyll content of semi-dominant *Oy1-N1989* mutants (Khangura *et al.* 2019). The *vey1* QTL was proposed to modulate the chlorophyll content of heterozygous *Oy1-N1989/+* mutants via cis-regulatory differences resulting in differential accumulation of the product encoded by the wild-type allele at *oy1*. The Mo17 allele at *vey1* (*vey1^Mo17^*) was associated with lower abundance of OY1 transcripts, whereas the B73 allele at *vey1* (*vey1^B73^*) is associated with higher accumulation of OY1. The effect of *vey1* on chlorophyll content is only visible in the presence of *Oy1-N1989*, indicating that this natural variant has a cryptic effect on the function of the MgChl complex.

In this study, we used controlled crosses to segregate *Oy1-N1989* and the modifier alleles at *vey1* (*vey1^B73^* and *vey1^Mo17^*) to generate populations of maize with a range of chlorophyll contents. We used this variation in chlorophyll to explore the effects of chlorophyll content on net CO_2_ assimilation. These changes in chlorophyll accumulation resulted in changes in net CO_2_ assimilation and photosynthetically-fixed carbon accumulation. Remarkably, we noticed that flowering time across material with differing photosynthetic rates and chlorophyll contents was dramatically different. We observed that reduced chlorophyll accumulation was associated with a delay in flowering time. Similar to the cryptic effects of *vey1* on chlorophyll content, the modifier allele had no effect on flowering time in the absence of the *Oy1-N1989* mutant allele. Chlorophyll content was consistently associated with earlier flowering and partial rescue of chlorophyll accumulation in the *Oy1-N1989* mutant by the *vey1* QTL accelerated flowering in the mutants but had no effect on wild-type siblings. In addition to measurements of net CO_2_ assimilation, the premature senescence of maize leaves, induced by sugar accumulation following sink removal, was also reduced by *Oy1-N1989* and further reduced by *vey1* alleles that decrease chlorophyll content and net CO2 assimilation. The effect of reduced photosynthate accumulation on flowering time was not specific to *Oy1-N1989* as mechanical removal of leaves, to reduce plant leaf area, also delayed flowering time. Thus, all of our results are consistent with an integrative measure of carbon assimilation linking energy status and flowering time in maize.

## Materials and Methods

### Plant materials

Our previously described stock of the *Oy1-N1989* mutant allele is maintained in the B73 background and is propagated by crossing heterozygous mutants (*Oy1-N1989/+*) to wild-type siblings (Khangura *et al.* 2019). The B73 introgressed stock of *Oy1-N1989* was used for crossing to various mapping populations. A total of 216 intermated B73 × Mo17 population (IBM-RILs), and 251 synthetic 10 doubled haploid lines (Syn10-DH) were crossed as ear-parents with *Oy1-N1989/+*:B73 pollen. The pollen of *Oy1-N1989/+*:B73 plants were also crossed on to the ears of 35 B73-Mo17 near-isogenic lines (BM-NILs) for QTL validation. Tables S1-S3 contain the full list of IBM-RILs, Syn10-DH, and BM-NILs used to develop F_1_ hybrid populations.

**Table 1.**
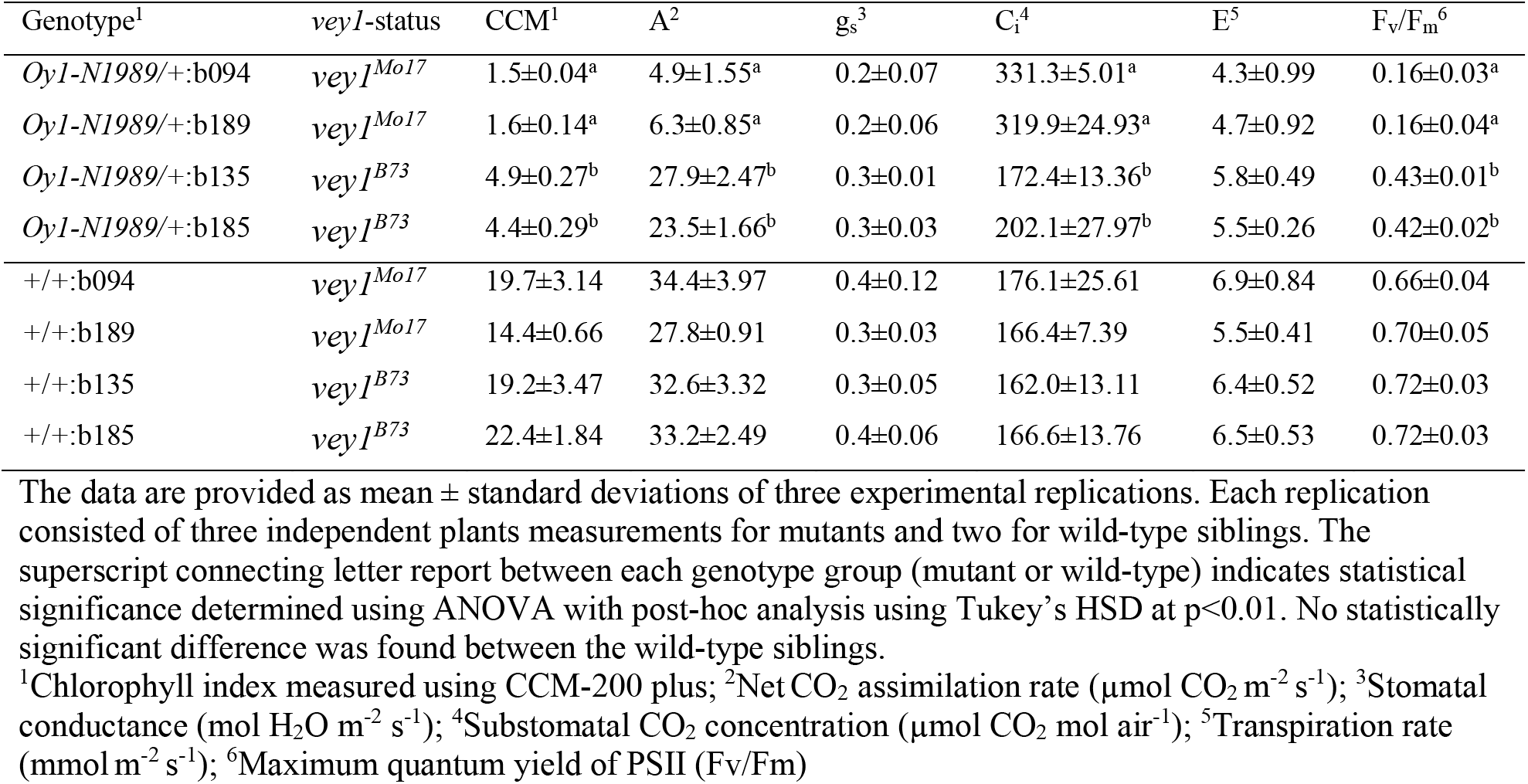
The chlorophyll index, gas-exchange, and maximum quantum yield of the mutant (*Oy1-N1989/+*) and wild-type (*+/+*) siblings in four independent B73-like NILs x *Oy1-N1989/+*:B73 F_1_ progenies.

### Field trials

All of the field experiments described in this study were conducted at the Purdue Agronomy Center for Research and Education (ACRE) in West Lafayette, Indiana. Each cross was evaluated as a single plot of 12-16 plants. Each plot derived from crosses with *Oy1-N1989/+*:B73 tester segregated for both mutant and wild-type siblings in approximately 1:1 ratio. Seeds were sown with a tractor-driven seed planter with plot length of 3.84 meters (m), alley length of 0.79 m, and inter-row spacing of 0.79 m. Standard crop management practices at Purdue in terms of fertilizer, pest, and weed control for growing field maize were adopted.

Each experiment was divided into blocks. Progenies from crosses of B73 and Mo17 ears to *Oy1-N1989/+*:B73 pollen were used as parental checks in each block. Parental checks were randomized within each block. The IBM-RILs F_1_ populations were evaluated as a single replication in 2013. Syn10-DH F_1_ populations were evaluated in 2016 with two full replications planted in randomized complete block design (RCBD). Hybrid progenies of *Oy1-N1989/+*:B73 with BM-NILs were screened in 2016 with five replications planted in RCBD.

### Field phenotyping and data collection

The mutant plants in each plot were identified visually as pale plants. The chlorophyll content in mutant and wild-type plants was approximated using a CCM-200 plus (Opti-Sciences Inc., Hudson, NH) as described in Khangura *et al.* 2019. We previously demonstrated correlation of 0.94 between CCM-200 plus values and chlorophyll *a*, chlorophyll *b*, and total chlorophyll contents (Khangura *et al.* 2019), and therefore used CCM-200 plus values (CCM) as a proxy for chlorophyll content in the materials described here. CCMI refers to measurements taken with the CCM-200 plus instrument from 25-30 days after sowing and CCMII refers to measurements taken 45-50 days after sowing. Mutants were tagged between the V5-V7 stages of development. Tagging is necessary as suppression of the *Oy1-N1989* mutant phenotype by *vey1* interferes with visual classification of mutant and wild-type siblings at maturity. Reproductive maturity in each F_1_ population was recorded separately on the wild-type and mutant plants. The date at which roughly half of the wild-type or mutant plants in a given plot were shedding pollen and had emerging silks from the primary ear was recorded as the date of anthesis and date of silking, respectively, for a given genotype. The dates of anthesis and silking for both wild-type and mutant genotypes were then subtracted from the date of planting to obtain respective wild-type or mutant days to anthesis (DTA), and days to silking (DTS). Further, the difference between DTA and DTS was used to derive the anthesis-silking interval (ASI); ASI = DTA-DTS. Wild-type and mutant trait values are denoted with a prefix WT and MT, respectively. Ratio and differences of these flowering time traits were also calculated as MT/WT and WT-MT, respectively.

A total of 15 F_1_ populations derived from B73-like NILs × *Oy1-N1989/+*:B73 cross were used to study induced leaf senescence. Seven of these B73-like NILs carried an introgression of *vey1* from Mo17 (*vey1^Mo17^*), whereas the other eight NILs had the B73 genotype at *vey1* (*vey1^B73^*). These NIL populations were planted in the field with at least two replications of each genotype planted in RCBD and two times separated by two weeks. The procedure for this experiment was adapted from Sekhon *et al.* 2012. Briefly, primary and secondary ears of both wild-type and mutant plants were covered with shoot bags before silk emergence. After 3-4 days of tassel shedding, roughly half of the shoot bags were removed, and these ears were allowed to open pollinate. Staggered rows of B73, in addition to the pollen shed within the row fully pollinated exposed ears. The day of shoot exposure was marked as 0 days after anthesis (DAA). Photographs were taken on the same date using plants from both planting dates to permit display of differences in the effect of DAA on phenotype severity.

### Genotypic and gene expression data

The genotypic data and other public datasets on various mapping populations used in this study have been described previously (Khangura *et al.* 2019). Briefly, the public marker dataset for IBM-RILs was obtained from MaizeGDB with 2,178 markers (Sen *et al.* 2010). The markers were reduced to 2,156 after removing duplicate variants, with ∼13.3 percent of missing data in the final dataset. Genotypic data consisting of 6611 SNPs for Syn10-DH lines was obtained from Liu et al. 2015. This dataset had no missing genotypes and was used as such for QTL analyses. Genotypes of the B73-Mo17 Near Isogenic Lines (BM-NILs) used for QTL validation were obtained from Eichten *et al.* 2011. Expression data of *oy1* locus in IBM-RILs were obtained from a public repository of the National Science Foundation grant (GEPR: Genomic Analyses of shoot meristem function in maize; NSF DBI-0820610; https://download.maizegdb.org/GeneFunction_and_Expression/ShootApicalMeristem/). This data consists of the expression of maize genes in the tissue derived from the shoot apex of 14 days old IBM-RILs seedlings. The expression data from each gene is normalized to reads per kilobase of transcript per million mapped reads (RPKM).

### Allele-specific expression (ASE) assay

Three replications of B73-Mo17 near-isogenic lines (BM-NILs) × *Oy1-N1989/+*:B73 F_1_ progenies were grown in the field. Mutant siblings derived from four B73-like NILs crossed with *Oy1-N1989/+*:B73 were selected for the ASE experiment. These NILs consisted of two B73-like NILs (b094 and b189) with *vey1^Mo17^*, and other two B73-like NILs (b135 and b185) with *vey1^B73^* genotype. Leaf tissue was harvested from the top fully-expanded leaf at the V3 developmental stage from the mutant siblings of the four B73-like NIL F_1_ plots. For each biological replicate, tissue was pooled from 4-5 seedlings to make one sample. The samples were stored at −80 °C until needed. The procedure of total RNA isolation, cDNA synthesis, and the ASE assay has been described previously in detail (Khangura *et al.* 2019). Briefly, one µg of DNase treated total RNA was used to synthesize cDNA. PCR was conducted using the forward oligonucleotide 5’-TCACCGTCTGCAATGTCGCCGCTC −3’ and reverse oligonucleotide 5’-AGTATGCCCCTGTTGGCCTTGGCG −3’ under following reaction conditions with 30 cycles of polymerization (94°C for 30s, 56°C for 30s, 72 °C for 30s and final extension for 2 minutes) to amplify the targeted region of OY1. The primer pair used in this assay flanked the SNP that causes the L176F amino acid substitution in the *Oy1-N1989* mutant allele. PCR products were sequenced on a MiSeq instrument (Illumina, San Diego, CA) at the Purdue Genomics Core Facility. Reads were aligned to a small reference sequence of B73 derived from targeted PCR region using the GATK packages (DePristo *et al.* 2011). Read counts derived from GATK was used to calculate allele-specific expression. Genomic DNA derived from B73 × Mo17 F_1_ hybrids resulted in roughly 1:1 read counts demonstrating no bias in the assay.

### Statistical analyses

Exploratory data analysis was done using JMP 13.0 (SAS Institute Inc. 2016). The pairwise correlations were calculated using the Pearson correlation coefficient. The average values of various traits from line-cross populations, IBM-RILs and Syn10-DH, were subjected to QTL analyses. QTL detection was done using a single interval mapping via the EM algorithm using the function “scanone” in R/qtl, a software package implemented in R (Broman *et al.* 2003; R core Team 2013). Similar results were obtained with composite interval mapping function “cim” in R/qtl (data not shown).

### Defoliation assay

The defoliation experiments were performed using maize inbred B73, sorghum (*Sorghum bicolor*) inbred BTx623, and green foxtail (*Setaria viridis*) inbred A10.1. These experiments were conducted in a greenhouse using mogul base high-pressure sodium lamps (1000 Watts) as the supplemental light source for L:D cycle of 16:8 hours, with the temperature set at 28°C (day-time) and 20°C (night-time). The maize inbred line B73 was defoliated at V3 leaf stage. All the leaves with a fully visible leaf collar were cut slightly above the ligule. Sorghum plants with three to four fully opened leaves (∼20 days after sowing in the greenhouse) were defoliated in a similar way. All fully expanded leaves at ∼15 days after planting, including those on tillers, were removed in *Setaria* plants. The time to reproductive maturity of both defoliated and undisturbed controls was recorded on maize as described above. For sorghum and *Setaria*, the date of head emergence, rather than anthesis, on every plant was recorded and deducted from the date of planting to obtain days to heading.

### Non-structural carbohydrate (NSC) quantification

The soluble sugars and starch were quantified from the mutant siblings of four B73-NILs × *Oy1-N1989/+*:B73 F_1_ population. Two of these NILs carried *vey1^Mo17^*, while the other two had *vey1^B73^* genotype. Plants were grown in the field with three replications using a RCBD. The top fully-expanded leaf at the V3 stage was harvested at 1:00 PM and transferred to liquid nitrogen. Leaf tissues were stored at −80°C until needed. Leaf samples were ground into a fine powder and ∼100 mg of the powder was used to extract sucrose, glucose, fructose, and starch using a previously described method (Leach and Braun 2016). The quantification of these NSC was done using a previously described method (Leach *et al.* 2017). Briefly, high-performance anion exchange (HPAE) chromatography (ICS-5000, Thermo-Fisher Scientific) was used to analyze the neutral fraction of the extract. Sugar standards were used to construct a standard curve, and samples were diluted to ensure that the detected values fell within the scope of this curve.

### Gas-exchange measurements

Gas-exchange measurements were taken on field-grown plants during the summer of 2017 at the Purdue ACRE farm. The gas-exchange measurements were taken on the third leaf on plants at the V3 stage between 11 AM and 1 PM using a LICOR LI-6400XT open photosynthesis system (LI-COR Inc., Lincoln, NE, USA). The F_1_ progenies used for these measurements consisted of four independent B73-NILs × *Oy1-N1989/+*:B73 cross, in which two NILs carried *vey1^Mo17^* introgression and the other two carried *vey1^B73^* allele. Plants were grown in a RCBD of three replicated blocks. For each genotype, nine plants were measured as three replicates in each of the three blocks. The following instrument conditions were maintained throughout the measurement period: an artificial light source with an intensity of 1700 µmol photosynthetically active radiation (PAR) m^-2^ s^-1^, air temperature of ∼31 °C, CO_2_ concentration of 400 mL L^-1^, air flow of 400 µmol s^-1^, and relative humidity of 50-60%. Leaf temperatures varied from 32-34°C during the measurements.

Chlorophyll fluorescence measurements were taken on the same leaves used for gas-exchange using a Handy PEA (Hansatech Instruments Ltd., Norfolk, UK). Leaves were dark-adapted for 20-30 min using leaf clips before taking measurements. The saturation pulse rate of 3000 µmol m^-2^ s^-1^ was used to measure the emission of chlorophyll fluorescence. The initial chlorophyll fluorescence yield (F_0_), the variable chlorophyll fluorescence yield (F_v_), and the maximum chlorophyll fluorescence yield (F_m_) were recorded. The maximum photochemical efficiency of PSII in dark-adapted leaves was obtained by calculating the ratio of F_v_/F_m_.

### Data availability

All phenotypic data from the QTL and NIL populations are attached to this manuscript as supplemental tables S1-S12 and available via figshare. All marker data was previously used in Khangura *et al.* 2019 and made available to the public via figshare (https://doi.org/10.25387/G3.7370948). All the seed stocks described in this study are available upon request.

## Results

### Negative effect of *Oy1-N1989* on time to reproductive maturity is exacerbated by Mo17

While preparing the material for our previous study (Khangura *et al.* 2019), we noticed that *Oy1-N1989* exhibited a consistent delay in flowering, as measured by the days to silking and days to pollen anthesis. Heterozygous *Oy1-N1989* mutant plants in the Mo17 × B73 hybrid genetic background flower up to two-weeks later than wild-type siblings (Figure 1; Table S4). The *Oy1-N1989* mutants also flower later in an isogenic inbred B73 background; however, the delay is only four days. By contrast, wild-type B73 × Mo17 F_1_ hybrid plants flower earlier than the wild-type B73 inbred plants. Maize is protandrous and tassels mature earlier than the ear-inflorescence. The effect of *Oy1-N1989* and flowering time was similar for both anthesis and silk emergence. The window of difference in maturity of the tassel and ear inflorescence, measured as anthesis-silking interval (ASI), is used as an indicator of plant stress in maize (Bolanos and Edmeades 1996). The ASI was wider in the wild-type siblings compared to the mutants in Mo17 × B73 hybrid background (Figure 1G) and not discernably different in the B73 inbred background. Thus, the delay in flowering does not seem to be due to a generic stress effect due to lower chlorophyll contents.

**Figure 1.**
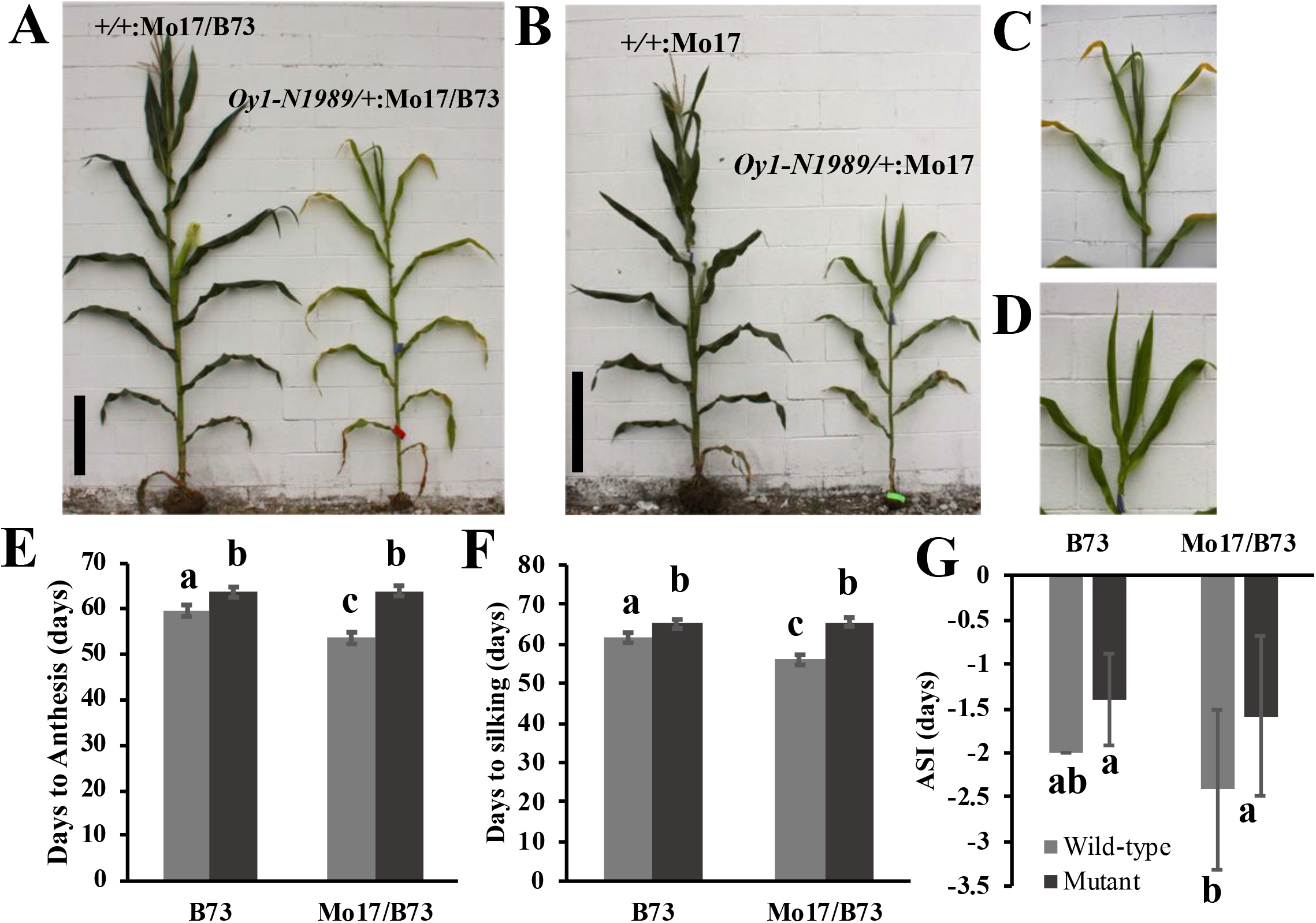
The reproductive maturity is delayed by *Oy1-N1989* allele in maize. The representative (A) wild-type (left) and mutant (right) sibling from Mo17 x *Oy1-N1989/+*:B73 cross (reproduced from Figure 1b in Khangura *et*. *al*. 2019), black scale bar = 50 cm. The representative (B) wild-type (left) and mutant (right) sibling from Mo17 x *Oy1-N1989/+*:Mo17 cross (BC_7_). The close-up view of the emerging tassel of mutant siblings (C) in panel A, and (D) panel B. The distribution of (E) days to anthesis, (F) days to silking, and (G) anthesis-silking interval (ASI) in the wild-type and mutant siblings in B73 and Mo17 x B73 (Mo17/B73) hybrid genetic backgrounds.

### Delayed reproductive maturity of *Oy1-N1989* mutants in B73 × Mo17 mapping populations maps to *vey1*

If the effect of genetic background on flowering time in *Oy1-N1989* mutants is due to variation in the accumulation of chlorophyll, we expect that the previously described *vey1* QTL from Mo17 should make this more severe (Khangura *et al.* 2019). To identify the genetic basis of the flowering time variation in *Oy1-N1989* mutants and test the effect of *vey1*, we recorded flowering time in wild-type and mutant F_1_ progenies from the crosses between *Oy1-N1989/+*:B73 pollen-parent with IBM-RILs and Syn10-DH lines. Hereafter, the hybrid populations developed from these crosses will be referred to as the IBM-RILs F_1_ and Syn10-DH F_1_ populations (Figure 2).

**Figure 2.**
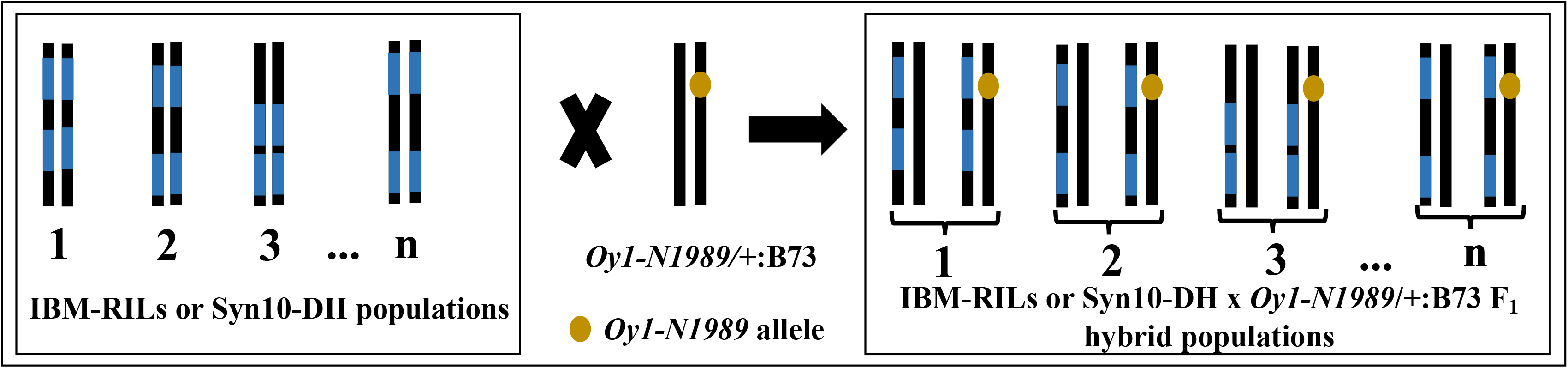
The schematic of crossing strategy used to map *Oy1-N1989* enhancer/suppressor loci using IBM-RILs (n=216) and Syn10-DH (n=251) populations. Black and blue colors indicate B73 and Mo17 genotypes. Chromosome 10 of the heterozygous pollen-parent *Oy1-N1989/+*:B73 is shown with a golden spot indicating *Oy1-N1989* mutant allele. The resulting F_1_ progenies from these crosses are depicted with state of chromosome 10 for each F_1_ testcross showing segregation of wild-type and mutant (with the golden spot) siblings.

Pairwise correlations were calculated between previously reported chlorophyll index measures (Khangura *et al.* 2019) and flowering time traits collected from the same plots (Tables S5 and S6). The chlorophyll index was measured at two time points CCMI (25-30 days after sowing) and CCMII (45-50 days after sowing). In the IBM-RILs crosses, wild-type CCMII displayed a weak but significant negative correlation with wild-type DTA and DTS. Similarly, in the Syn10-DH crosses wild-type CCMI displayed a significant weak negative correlation with wild-type DTA and DTS. This indicates that the phenomena observed in our mutants, reduced chlorophyll content associated with a delay in flowering time was true in the wild-type populations as well, but much less obvious. The variation in chlorophyll content in wild-type plants was not predictive of either mutant CCM or flowering time in the mutants (Figure 3, Table S5, and S6) indicating that the variation in CCM was not under the same control in the mutant and wild-type subpopulations. A dramatic and obvious negative correlation was observed between CCM trait values (CCMI and CCMII) and flowering time in the mutant siblings in both IBM-RILs and Syn10-DH F_1_ populations. As was observed in the *Oy1-N1989/+* B73 inbred stock and B73 × Mo17 hybrids *Oy1-N1989/+* mutants, mutants in these test-cross populations also showed a clear increase in mean values for days to anthesis and silking compared to wild-type siblings (Figure 4).

The frequency distribution plot of days to anthesis in mutant siblings of IBM-RILs and Syn10-DH F_1_ populations displayed a bimodal distribution, suggesting a single polymorphic locus segregating between B73 and Mo17 was the basis of flowering time variation in *Oy1-N1989* mutants (Figure 5). QTL mapping detected a single QTL on chromosome 10 at similar linkage positions for all mutant derived flowering traits (Figure 5; Tables S7 and S8). This corresponds to the *vey1* locus that we previously described as a major-effect QTL that controls chlorophyll biosynthesis only in the presence of the *Oy1*-*N1989* allele (Khangura *et al.* 2019). QTL mapping for various direct and derived mutant flowering time traits such as days to flower in the mutant heterozygotes (MT_DTA and MT_DTS), ratio of days to flower derived from mutant and wild-type siblings (Ratio_DTA and Ratio_DTS), and the difference in days to flower between the wild-type and mutant siblings (Diff_DTA and Diff_DTS) all detected *vey1* in both mapping populations and exceeding permutation-estimated significance thresholds (alpha<0.05). The *vey1* QTL explained ∼40-48% of the phenotypic variation for these flowering traits in the IBM-RIL crosses, and >65% variation in Syn10-DH F_1_ crosses. As expected, the *Oy1-N1989* enhancing *vey1^Mo17^* allele was associated with a delay in flowering time in the mutant hybrid siblings (Tables S7 and S8). In addition to *vey1*, an additional QTL controlling both mutant anthesis and silking was detected in the Syn10-DH F_1_ population on chromosome 2 which we call *other oil yellow1 flowering time locus1* (*oof1*). This QTL explained ∼7-8% variation in flowering time with the Mo17 allele at this locus resulting in a delay of 2-3 days in reproductive maturity of mutant siblings.

**Figure 3.**
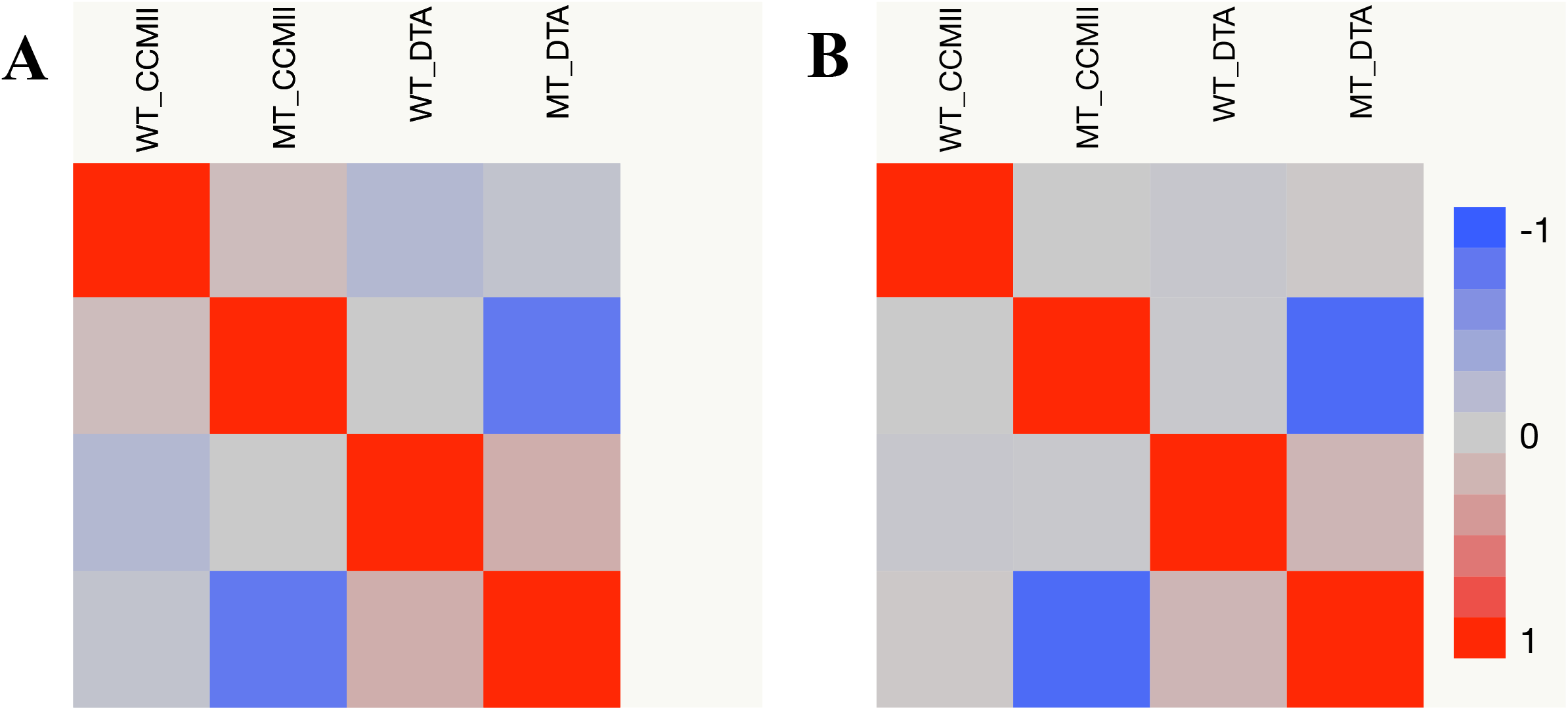
The pairwise correlation matrix showing color-coded correlation values between wild-type CCMII (WT_CCMII), mutant CCMII (MT_CCMII), wild-type days to anthesis (WT_DTA), and mutant days to anthesis (MT_DTA) in (A) IBM-RILs x *Oy1-N1989/+*:B73, and (B) Syn10-DH x *Oy1-N1989/+*:B73 F_1_ test cross populations.

**Figure 4.**
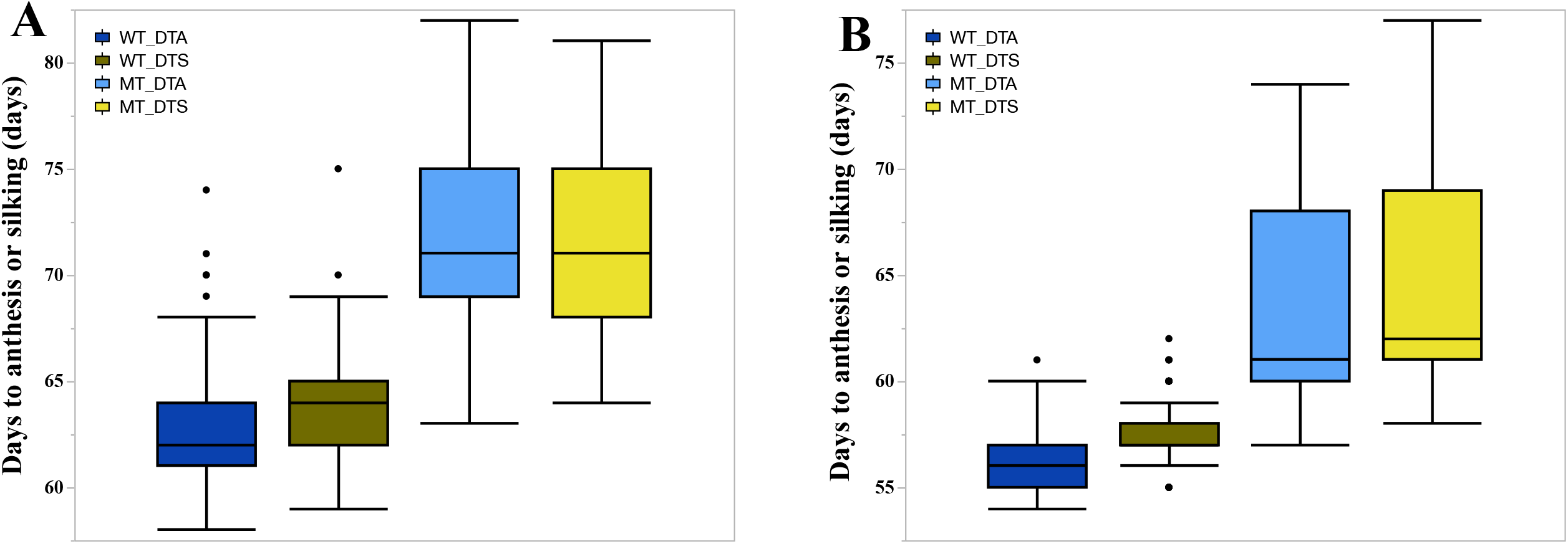
The distribution of days to flower (anthesis and silking) in the wild-type and mutant siblings in (A) IBM-RILs x *Oy1-N1989/+*:B73, and (B) Syn10-DH x *Oy1-N1989/+*:B73 F_1_ test cross populations. Abbreviations: wild-type days to anthesis (WT_DTA), wild-type days to silking (WT_DTS), mutant days to anthesis (MT_DTA), and mutant days to silking (DTS).

**Figure 5.**
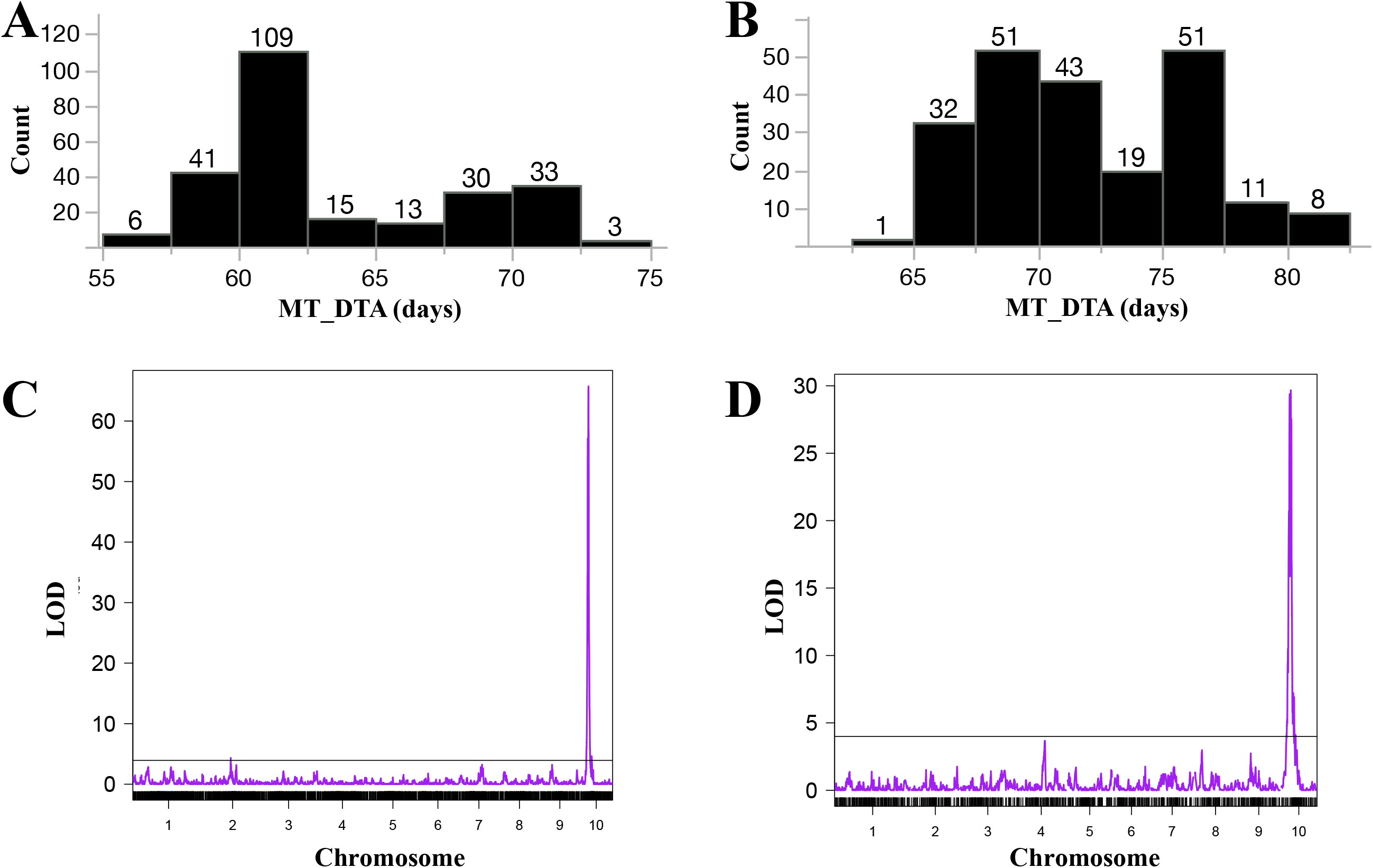
The distribution of days to 50% anthesis of the mutant siblings (MT_DTA) in (A) Syn10-DH x *Oy1-N1989/+*:B73, and (B) IBM-RILs x *Oy1-N1989/+*:B73 F_1_ population. Genome-wide QTL plot of MT_DTA in (C) Syn10-DH, and (D) IBM-RILs F_1_ population. The x-axis in (C) and (D) indicates the chromosome number, and y-axis indicates the logarithm of odds (LOD) of tested markers. Black horizontal bar indicates permutation testing based threshold for QTL detection.

Because of the very large effect of *vey1*, and the mild segregation distortion at this locus in both bi-parental populations (Khangura *et al.* 2019), weak QTL might be detected by spurious linkage between chr2 and chr10 markers. To test this, we carried out multiple regressions using top markers at *oof1* and *vey1* as independent variables and mutant DTA in the Syn10-DH as a dependent variable (data not shown). In this analysis, both *vey1* and *oof1* remained significant factors in the multiple regression and explained 62% and 2% of the variance in flowering time, respectively. Inclusion of a *vey1* × *oof1* interaction term did not improve the fit of the model, did not eliminate the significance of the *oof1* term, and the interaction term was not a significant variable. Moreover, we did not detect additional QTL by including these as covariates in an additional genome-wide scan. Therefore, we propose that *oof1* is a novel QTL contingent upon the *Oy1-N1989* mutation and genetically independent of *vey1*.

The ASI of mutant and wild-type siblings in the test cross populations were not discernably different, just as we observed in the mutant parents. QTL mapping for this trait did not detect any loci controlling ASI in either the mutants or the wild type siblings. In addition, we did not detect any QTL for flowering time in wild-type siblings (Tables S7 and S8). Thus, *Oy1*-*N1989* was epistatic to both *vey1* and *oof1* QTLs suggesting a role for each locus in controlling either photosynthesis or chlorophyll metabolites in the regulation of flowering time.

We further validated the effect of the *vey1* critical region using a set of NIL that vary at the *vey1* QTL from the recurrent background. F_1_ progeny of these NIL and *Oy1-N1989*/+ produced matched wild-type and *Oy1-N1989*/+ heterozygous mutant NIL F_1_ hybrids. The *vey1^Mo17^* allele delayed flowering time in *Oy1-N1989/+* mutants crossed to both B73 and Mo17 recurrent backgrounds when compared to NILs carrying *vey1*^B73^ (Figure 6, Figure S3, and Table S3). No effect of *vey1* introgression from either parent was visible on the flowering traits of the wild-type siblings. The reproductive maturity of mutant B73-like NILs carrying *vey1^Mo17^*, and reciprocal introgression of *vey1^B73^* into Mo17 background displayed the opposite effect on flowering time in the F_1_ mutants (Figure 6). Thus, these results clearly show the single locus effect of *vey1* on flowering time in maize in *Oy1-N1989*-contingent manner in both isogenic inbred (B73) and hybrid (Mo17 × B73) background.

**Figure 6.**
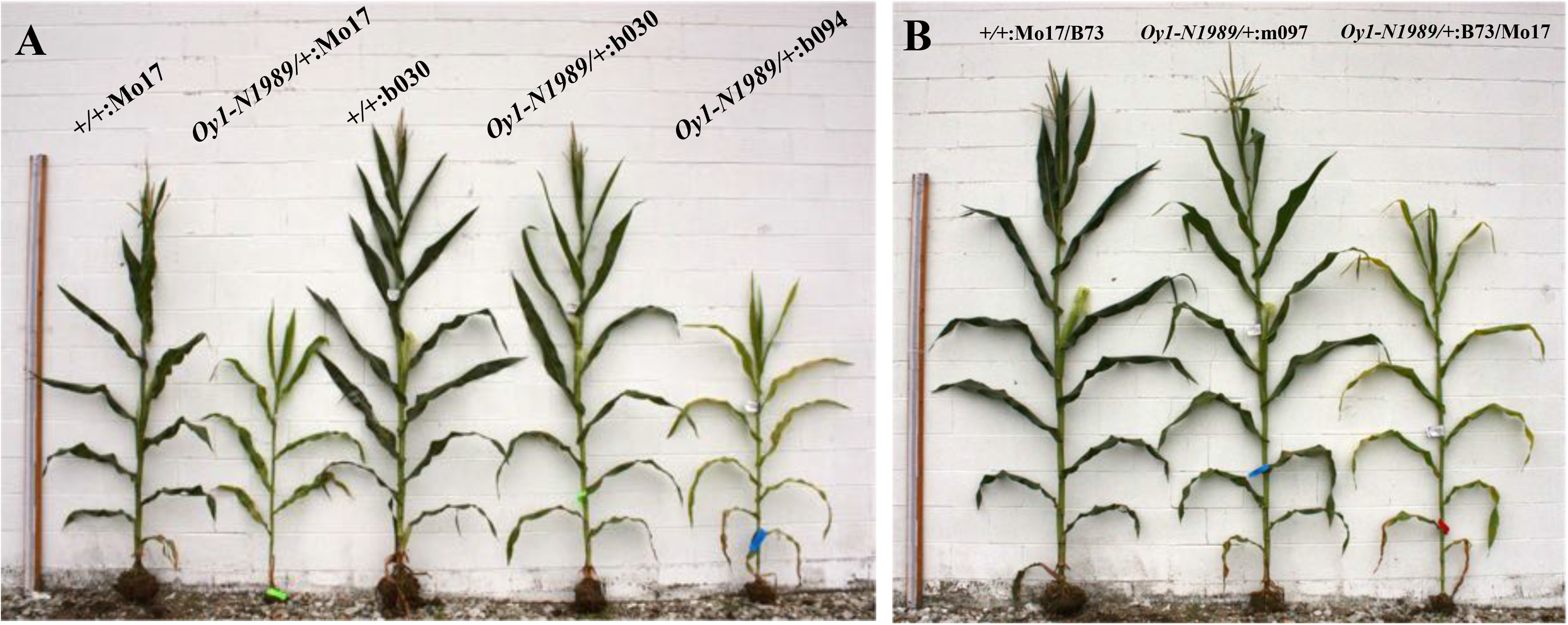
Effect of *vey1* locus from B73 and Mo17 on flowering time in reciprocal isogenic backgrounds. Representative plants showing Delayed maturity of the heterozygous mutants in isogenic Mo17 (BC_7_ generation) inbred background, compared to the wild-type siblings. Mutant and wild-type siblings in b030 (B73-like NIL with *vey1^B73^*) shows early flowering while the isogenic B73 introgression of the *vey1^Mo17^* allele in b094 NIL exhibits delayed flowering. (B) Early maturity in the heterozygous mutant in m097 (Mo17-like NIL with *vey1^B73^*) compared to the mutant in B73 x Mo17 F_1_ hybrid background. Measuring stick on the left in both panels is 243 cm.

### Expression polymorphism in B73-like NILs is consistent with cis-acting regulatory polymorphism at *vey1*

Our previous study looking at the suppression of *Oy1-N1989* mutant phenotype using chlorophyll accumulation identified a cis-eQTL at *oy1* in the IBM-RILs population (Khangura *et al.* 2019). Normalized expression (expressed as RPKM) of OY1 derived from 14 days old shoot apices of IBM-RILs (Li et al., 2013; 2018) were used for this analysis. The top marker, isu085b, used in the detection of this cis-eQTL was also one of the top significant markers for mutant flowering time traits (Figure S1 and Table S9). Regression of OY1 expression and flowering traits collected in IBM-RILs F_1_ population identified a significant linear relationship between gene expression in wild-type inbred lines and flowering time measurements from mutant F_1_ siblings (Figure S2 and Table S9). Roughly 21% of the variation in mutant DTA could be explained by OY1 expression in the IBM-RILs shoot apices. OY1 expression did not predict any variation in wild-type DTA.

The allele specific-expression (ASE) assay in our previous study identified bias in expression with the wild-type *oy1* allele from Mo17 displaying lower expression than the wild-type *oy1* allele from B73 (Khangura *et al.* 2019). Our previous ASE work compared mutant plants in two different genetic backgrounds (inbred vs hybrid) which can complicate the interpretations. To overcome this limitation, an ASE assay was designed to test bias at *oy1* using near-isogenic lines in the B73 background. These B73-like NILs consisted of two independent NILs with *vey1^Mo17^* introgression, and two independent NILs with B73 genotype at *vey1*. Consistent with the previous ASE results, a significantly greater proportion of expression was derived from the *Oy1-N1989* mutant allele when the wild-type *oy1* allele was contributed by *vey1^Mo17^* introgression as compared to the isogenic mutant siblings carrying the wild-type *oy1* allele from *vey1^B73^* introgression (Table S10).

### Net CO_2_ assimilation and sugar metabolism is reduced in *Oy1-N1989* mutants in *vey1*-dependent manner

We measured net CO_2_ assimilation, sub-stomatal CO_2_, photosystem II fluorescence, and photosynthate accumulation in enhanced and suppressed *Oy1-N1989/+* mutants. A previous study in maize has shown that reduction in chlorophyll levels in the leaves leads to a reduction in photosynthetic rate (Huang *et al.* 2009). A similar reduction in photosynthetic rate should be displayed by *Oy1-N1989* mutants and the Mo17 allele should show a greater reduction in photosynthesis compared to the B73 allele. We tested this using F_1_ progenies derived from the same four B73-like NILs used for the ASE experiment. The negative effect of *vey1^Mo17^* introgression on chlorophyll accumulation was visible in the *Oy1-N1989* mutant allele background (Figure 6). As expected, photosynthetic rate (A) was reduced in mutants as compared to wild-type siblings, and mutants were modified further by the *vey1* genotype (Table 1). Photosystem II efficiency (Fv/Fm) measurements indicated higher photo-oxidative damage to the photosystem in enhanced mutant plants compared to the suppressed mutants. Wild-type siblings of all four B73-like NIL F_1_ progenies showed no statistically significant difference for chlorophyll and gas-exchange measurements. This indicates that, just as for chlorophyll content, the *vey1* QTL affects net CO_2_ assimilation in the presence of the *Oy1-N1989* mutant allele. No differences in stomatal conductance (g_s_) or transpiration (E) were observed in the mutant plants, however, severe mutant plants showed significantly higher accumulation of intracellular CO_2_ (Ci) compared to the suppressed mutants suggesting the failure of the enhanced mutant plants to uptake CO_2_ (Table 1).

These differences in photosynthetic rate should result in a decrease of non-structural carbohydrate accumulation in the photosynthetic leaf tissue. The levels of non-structural carbohydrates were determined from leaves of the same mutant plants that were used for gas-exchange measurements. Levels of sucrose, glucose, fructose, and starch were measured in these samples. All of these showed a significant reduction in the enhanced mutants compared to the suppressed mutants in the B73 isogenic background (Figure 7 and Table S11). Lower levels of sugars and starch in the *Oy1-N1989/+* mutant heterozygotes enhanced by a *vey1^Mo17^* allele is consistent with the observation of lower chlorophyll levels and photosynthetic rates in these genotypes compared to the suppressed mutant NILs.

**Figure 7.**
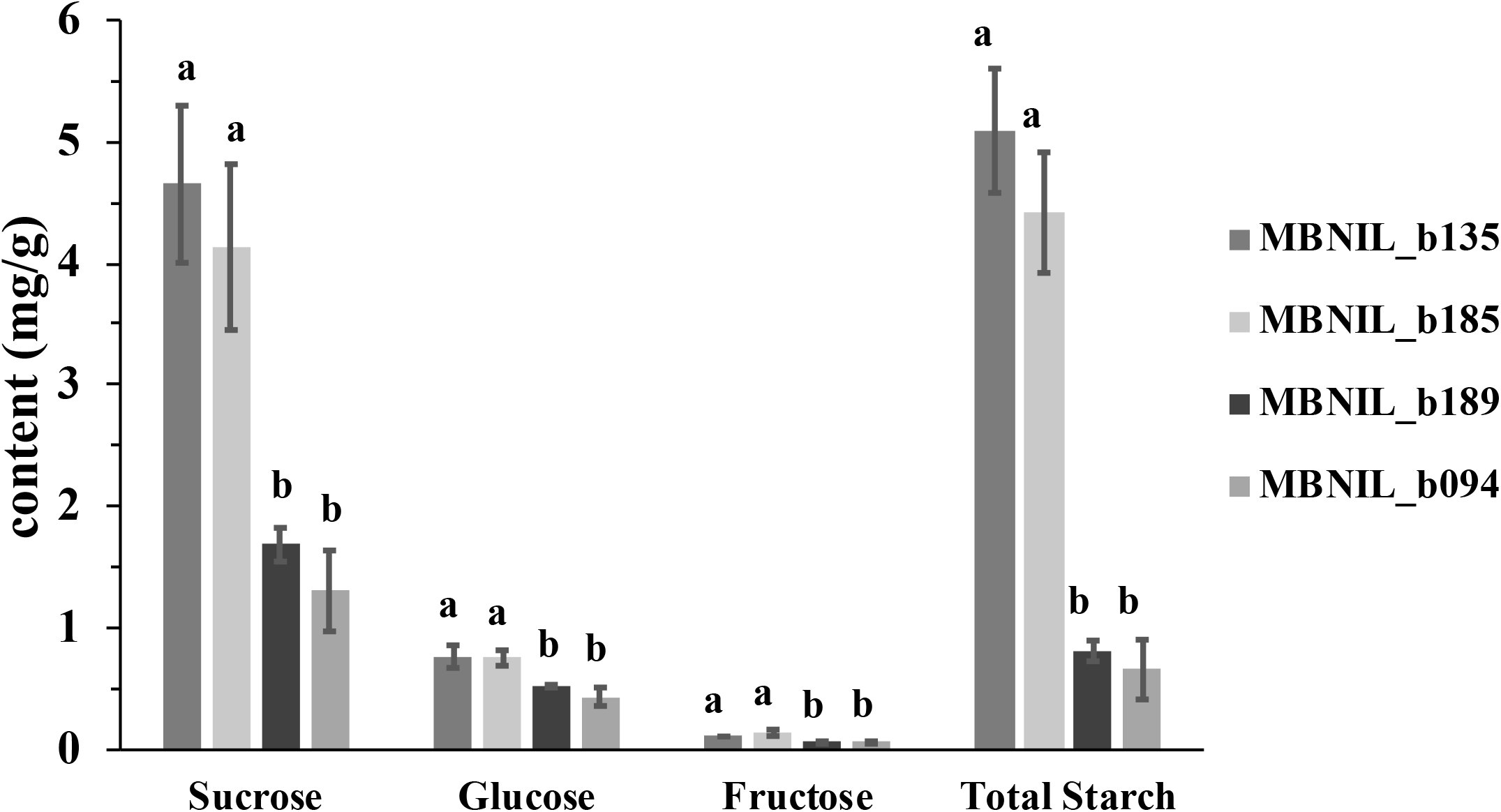
The effect of alleles at *vey1* on leaf soluble sugars and starch content in *Oy1-N1989/+* heterozygotes in the isogenic B73 background. B73-like NILs b135 and b185 have *vey1^B73^* introgression, whereas b189 and b094 have *vey1^Mo17^* introgression. The values of different sugars and starch are reported as mg/g of fresh weight.

### Defoliation of *Zea mays*, *Sorghum bicolor*, and *Setaria viridis* delays reproductive maturity

The existing literature suggests sugars and carbohydrate metabolism play an important role in regulating flowering time in plants (Ohto *et al.* 2001; Seo *et al.* 2011; Wahl *et al.* 2013). We hypothesized that removal of source tissue should mimic the sugar starvation observed in *Oy1-N1989* mutants. We used mechanical defoliation to reduce photosynthetic surplus of the plant. The choice of this treatment was intended to deprive plants of leaf area and photosynthate early in development to separate the block in chlorophyll biosynthesis and the loss of photosynthate, which are coupled in the *Oy1-N1989* mutant study. We conducted this experiment using wild-type inbred strains of maize and two other monocot species: *Sorghum bicolor* (sorghum) and *Setaria viridis* (green foxtail). Each species was defoliated at an early vegetative stage when only few leaves had fully expanded. Defoliation delayed flowering in all three species (Figure 8). Maize, sorghum, and green foxtail displayed a delay in flowering by about 13, 7, and 4 days, respectively, compared to the control plants. Remarkably, one-week post-defoliation, newly-emerged leaves displayed a pale leaf color. Chlorophyll estimation using CCM found a reduction in the leaf greenness in the defoliated treatments compared to the control samples (Figure 8). The newly-emerged leaves of defoliated maize, sorghum, and green foxtail showed ∼35%, ∼48%, and ∼58% reduction in CCM, respectively, compared to control plants. Subsequent leaves emerging from the defoliated plants displayed normal leaf color suggesting recovery of the plants from defoliation. We propose that early season defoliation results in the removal of source tissue that might be critical for vegetative to floral transition in grasses but a direct effect of chlorophyll accumulation cannot be ruled out.

**Figure 8.**
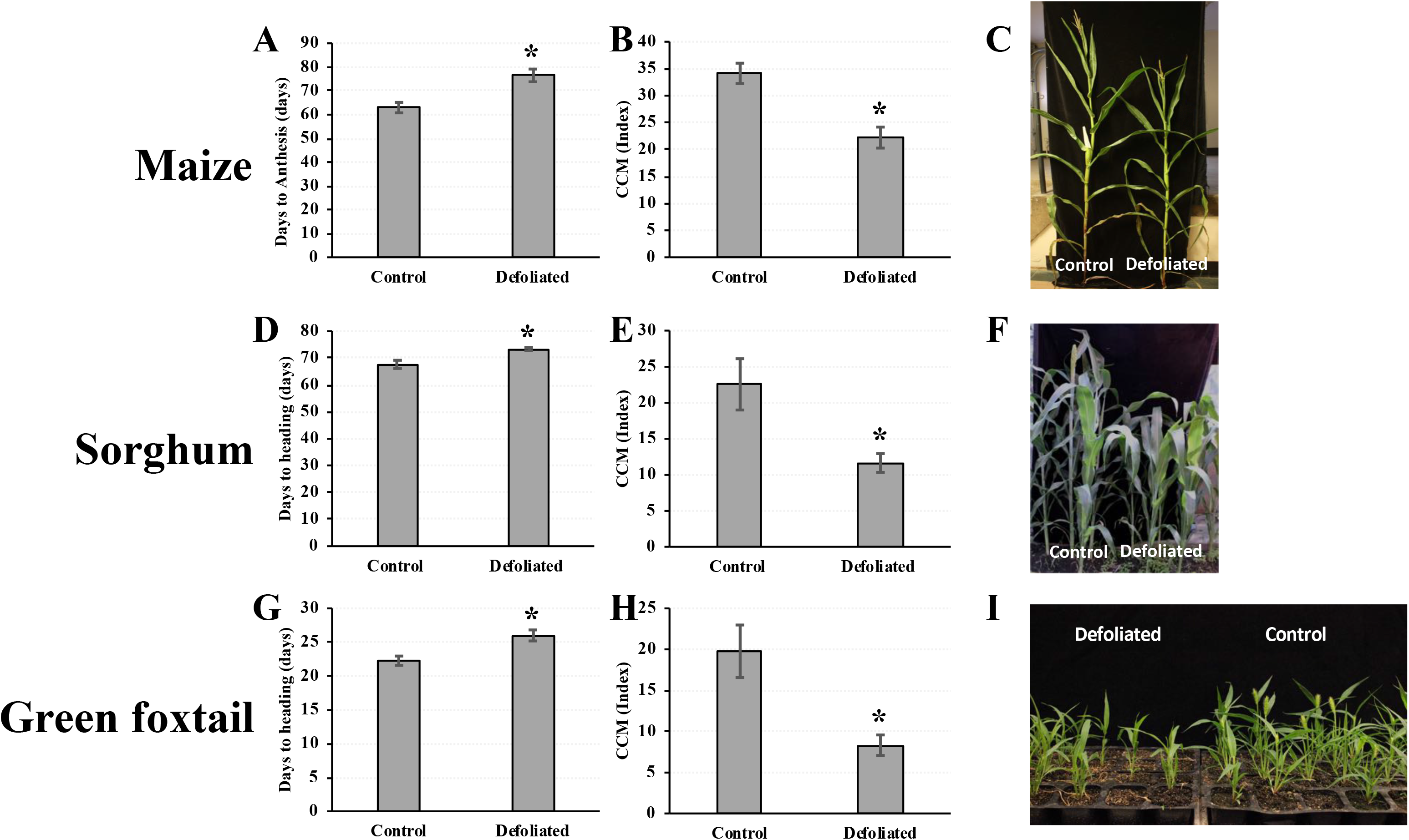
The effect of defoliation on flowering time and chlorophyll in the newly emerged leaves of (A-C) maize, (D-F) sorghum, and (G-I) green foxtail. The asterisk indicates significant difference between treatment means using student’s t-test at p<0.05.

### Leaf senescence is suppressed by *Oy1-N1989* mutants in a *vey1*-dependent manner

Leaf senescence can be induced in maize by sucrose accumulation in the leaves. This can be accomplished genetically by disrupting sucrose transport (Braun *et al.* 2006; Baker and Braun 2008; Slewinski *et al.* 2009) or by preventing the maize ears from acting as a sink (Allison and Weinmann 1970; Sekhon *et al.* 2012, 2019). Previous studies have shown that maize leaf senescence caused by pollination prevention or ear removal before pollination is genotype-dependent (Ceppi et al. 1987). We tested the effect of sugar accumulation in mutant B73-like NILs on induced leaf senescence by pollination prevention. Given the variation in leaf sugar in *Oy1-N1989/+* (Figure 7) we expect the mutants exhibit less or later senescence following pollination prevention and modulation of this effect by *vey1*. We observed that 30 days-after-anthesis (DAA), the top leaves of unpollinated wild-type B73 plants showed complete senescence with only a few green patches. Unpollinated *Oy1-N1989/+* F_1_ mutant plants crossed to the *vey1^B73^* NIL background were green and showed only a few patches of anthocyanin accumulation and cell death on the top leaves at 30 DAA (Figure 9 and Table S12). Consistent with the lower chlorophyll and NSC accumulation, unpollinated *Oy1-N1989/+* mutant F_1_ plants crossed to the B73 NIL background containing the *vey1^Mo17^* introgression did not show any sign of leaf senescence at 30 DAA. By 42 DAA, unpollinated *Oy1-N1989/+* mutants with the *vey1^B73^* allele and all unpollinated wild-type plants showed leaf senescence. Even at 42 DAA, the enhancement of *Oy1-N1989/+* by the *vey1^Mo17^* allele resulted in substantially less cell death and anthocyanin accumulation.

**Figure 9.**
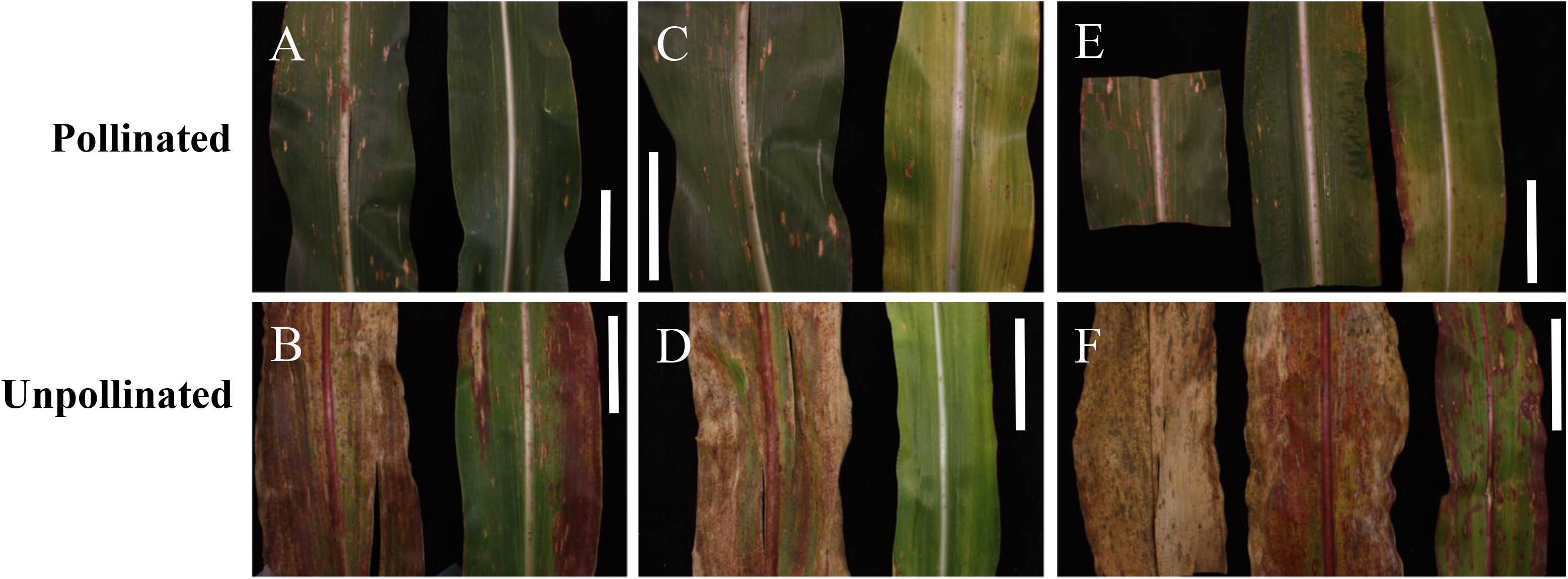
The effect of *vey1* on senescence induced by pollination prevention. Pictures of the representative primary ear leaf derived from plants at 30 and 40 days-after-anthesis (DAA) either with open pollinated (top panels: A, C, and E) or unpollinated (bottom panels: B, D, and F) ears. The representative primary ear leaf from (A-B) B73 wild-type (left) and mutant sibling (right) at 30DAA, (C-D) wild-type (left) and mutant (right) sibling from F_1_ between B73-like NIL-b107 (with *vey1^Mo17^* introgression) x *Oy1-N1989/+*:B73 at 30 DAA, (E-F) wild-type (B73-like NIL-b107), mutant B73-like NIL-b135 with *vey1^B73^* (middle), and mutant B73-like NIL-b107 with *vey1^Mo17^* allele (right) at 42 DAA. The scale bar in each figure is 6.35 cm.

## Discussion

Our previous work identified a modifier, *vey1*, that affects the chlorophyll accumulation in *Oy1-N1989/+* heterozygotes (Khangura *et al.* 2019). The *vey1* polymorphism(s) are common natural variant(s) linked to the *oy1* locus of maize. We proposed that *vey1* results from cis-acting regulatory polymorphisms that affect the expression of OY1. The suppression of the mutant, and accumulation of chlorophyll, follows the proportion of wild-type and mutant transcript levels (Table S10 and Khangura *et al.* 2019). In this work, we describe a delay in reproductive maturity in the *Oy1-N1989* mutants and demonstrate that *vey1* encodes a strong modifying QTL altering the flowering time of mutant siblings in all mapping populations (Figures 5 and 6; Tables S7 and S8). Just as the detection of *vey1* for CCM was contingent on the *Oy1-N1989* mutant allele in the background, there was no effect of the *vey1* genotype on flowering time in wild-type siblings. We observed the same marker, isu085b, had the strongest effect on chlorophyll content, OY1 transcript abundance in shoot apices (Khangura *et al.* 2019), and variation in flowering time of IBM-RILs × *Oy1-N1989/+* mutant F_1_ mutant siblings (Table S9 and Figure S2). Taking these observations together, we propose that the cis-acting eQTL at *oy1* is affecting chlorophyll level in *Oy1-N1989* mutants, and that the alteration in photosynthesis through perturbed chlorophyll metabolism affects flowering time variation in mutant siblings in these populations.

Our previous work on *vey1* has focused on the cryptic nature of the variation, and the interaction between the *vey1* QTL and the *Oy1-N1989* mutant allele. The experiments presented here also suggest chlorophyll content as a heretofore unappreciated correlate of flowering time in wild-type maize plants (Figure 3; Tables S5 and S6). We observed that CCM values in both test cross populations, were negatively correlated with wild-type days to reproductive maturity. This suggests a role for the determinants of variation in chlorophyll contents in regulation of flowering time in maize perhaps via changes to photosynthetic output or signaling. As no QTL were detected for CCM in the wild-type siblings in our previous work (Khangura *et al.* 2019), it also suggests that the mechanism responsible for the covariation between CCM and flowering time is independent of *vey1*. This is further strengthened by the absence of an effect of *vey1* on flowering time in the wild-type siblings in our mapping populations and NIL experiments (Table S3 and Figure S3). Further experiments are required to validate or reject a causal relationship between CCM and flowering time in wild-type siblings. Experiments using populations with greater recombination or allelic diversity, such as in an association panel, should disrupt most fortuitous linkage and would provide a second test of this phenotypic correlation and either argue for or against additional exploration of this relationship. The crosses of *Oy1-N1989/+* to the association panel analyzed for CCM in a previous study (Khangura *et al.* 2019) could be replanted and measured for flowering time of wild-type and *Oy1-N1989/+* mutant sibling pairs. Candidate gene testing of epistatic interactions between *Oy1-N1989*, *vey1*, and the known flowering time regulators segregating in that population (e.g. *zmmads69*, *cct10*, *zcn8*, *dlf1*, and *vgt1)* could provide some insight. Epistasis, indicating interaction between chlorophyll biosynthetic disruption and developmental determinants of the transition to flowering, would be consistent with photosynthesis acting as part of the autonomous pathway whereas no genetic interaction would be consistent with the slow growth of plant organs in a compromised background resulting in the observed reproductive delays.

Even though the Syn10-DH and IBM-RILs are derived from the same parents, they differ in the method of development and rates of recombination (Liu et al. 2015; Hussain et al. 2007; Lee et al. 2002). Thus, these two populations yield different levels of resolution for QTL detection. Our previous analysis to fine map *vey1* QTL using CCM values showed Syn10-DH to have a higher mapping resolution than the IBM-RILs population (Khangura *et al.* 2019). We also observed different QTL for flowering time in these populations. QTL analysis in Syn10-DH F_1_population detected the *oof1* QTL on chromosome 2 affecting MT_DTA and MT_DTS as well as the major-effect locus *vey1* (Table S8). The detection of *oof1* was dependent on the presence of the *Oy1-N1989* mutant, but was neither contingent nor displayed any epistatic interactions with *vey1*. Thus, *oof1* appears to be a novel locus of independent mechanism affecting flowering time in the *Oy1-N1989* mutants. This QTL on chromosome 2 was not detected for any other trait in the Syn10-DH and IBM-RILs testcross populations with *Oy1-N1989/+*:B73 in the current study nor was it identified as a modifier of chlorophyll content in our previous study (Khangura *et al.* 2019). The amount of phenotypic variation explained by *vey1* in flowering time was higher in the Syn10-DH experimental material than the IBM-RILs testcross population by ∼20% (Tables S7 and S8). Greater variation in flowering time in the IBM-RILs testcross progenies that could not be modeled by marker genotypes may result from higher residual heterozygosity or greater rates of pollen contamination during the single seed descent (SSD) procedure used to generate the IBM-RILs (Lee et al. 2002). Heterozygosity is expected to be negligible in Syn10-DH population because of the DH procedure employed to fix allele segregation during population development (Hussain *et al.* 2007). In addition, we had a moderate increase in sample size in the Syn10-DH (251 lines) as compared to the IBM-RILs (216 lines) that is expected to result in a marginally greater power to detect QTL in the Syn10-DH F_1_ populations.

Chlorophyll levels are correlated with the rate of photosynthesis in plants (Huang *et al.* 2009). Some controversy has been reported in mutants of soybean affected in an ortholog of *oy1* with some reports showing little impact on photosynthesis (Sakowska *et al.* 2018) and others clearly demonstrating a linear relationship between chlorophyll variation and photosynthesis (Walker *et al.* 2017). The reasons for conflicting conclusions results from soybean mutants are not fully clear but it warrants some caution in making simplistic interpretations about the impact of chlorophyll deficient mutants on photosynthesis. In our study, net CO_2_ assimilation was associated with the severity of the *Oy1-N1989* mutant phenotype. The effect of chlorophyll loss on CO_2_ assimilation measurements using a LICOR instrument was somewhat non-linear. A nearly 5-fold reduction in CCM in *Oy1-N1989* NILs carrying a B73 allele at *vey1* resulted in in a 22% reduction in net CO_2_ assimilation while the 10-fold reduction in CCM in *Oy1-N1989* NILs carrying a Mo17 allele at *vey1* resulted in a 64% reduction in net photosynthetic rate, compared to their isogenic wild-type siblings (Table 1). Ultimately, both reductions resulted in less accumulation of free sugars and starch, with a substantially greater reduction in the enhanced mutant NILs (Figure 7; Tables 1 and S11).

The allelic interactions of *Oy1-N1989* and wild-type *oy1* alleles are consistent with the inductive role of carbohydrate status on floral transition (Ohto et al. 2001; Seo et al. 2011; Wahl et al. 2013; Minow et al. 2018). Additional work exploring proposed carbohydrate signaling metabolites, such as T6P and organic acids, and analyses of the downstream floral integrators such as the FT orthologs of maize are still needed to link our results to the existing models of floral transition regulation in maize (Minow *et al.* 2018).

One complexity that we observed in our data is that defoliation did not separate photosynthate levels from changes in chlorophyll (Figure 8). The mechanical removal of leaves resulted in changes to CCM in the newly emerging leaves of defoliated plants. As a result, a mechanical treatment served to highlight the interconnected nature of plant metabolism: large changes to any feature of central metabolism or plant physiology results in large changes to all of central metabolism and plant physiology. In an effort to separate photosynthate and chlorophyll accummulation, one can conceive of alternate experiments such as measuring the time to maturity in different light intensities achieved using neutral shade cloth to produce a gradient of photosynthetically active radiation and sugar accumulation. Similarly, plants could be sprayed with low doses of chemicals such as 3-(3,4-dichlorophenyl)-1,1-dimethylurea (DCMU) that disrupt electron transfer during the light reactions to reduce photosynthetic output of plants without altering light fluence experienced by other photoreceptors. However, all these experiments may suffer from the same confounding of NSC and chlorophyll as the defoliation experiment. As a result, researchers should quantify chlorophyll after these treatments as any change in chlorophyll complicate our ability to uncouple chlorophyll metabolism and sugar accumulation as demonstrated by our defoliation experiment and all the carbohydrate partitioning mutants of maize studied to date.

Two other subunits of magnesium chelatase are encoded by genes that were identified in Arabidopsis as *genomes uncoupled* mutants, *gun4* and *gun5*, with altered retrograde, plastid-to-nuclear, signaling with defects in chlorophyll metabolism (Susek *et al.* 1993; Mochizuki *et al.* 2001; Larkin *et al.* 2003). Multiple studies have demonstrated retrograde signaling mutants with defects in chlorophyll metabolism (Hernández-Verdeja and Strand 2018) and circadian rhythm (Jones 2019). Retrograde signaling events are also associated with cell death, chloroplast development, and etiolation but it acts via a number of modifiers of flowering, including phytochromes, phytochrome signaling components, blue light perception via *crytochrome1*, and circadian rhythmicity of key genes that affect flowering time. While retrograde signaling is an attractive alternative model to sugar signaling for the phenomena reported here, mutants in orthologs of *oy1* in both monocots and dicots (Mochizuki *et al.* 2001; Gadjieva *et al.* 2005) have been tested for a *genomes uncoupled* phenotype and they did not perturb retrograde signaling. This makes it unlikely that *Oy1-N1989* and *vey1* are altering flowering time via aberrant retrograde signals. Additional experiments are necessary to clearly separate the effects of chlorophyll metabolite levels from photosynthate levels to independently test their effects on flowering time in maize.

We have known that defoliation is practiced in some maize nurseries to stagger flowering time and permit intercrossing of lines with divergent reproductive maturities. We had presumed that this was based on published research. Remarkably, we were not able to find a reference for this practice. Previous work on defoliation in maize has looked at the effect of early and late season defoliation on growth, and yield components but did not report flowering times (Crookston and Hicks 1988; Pearson and Fletcher 2009). As a result, Figure 8 provides information that was informally shared within the maize genetics research community but not described formally. In addition, we extended this observation from maize to both sorghum and green foxtail, indicating that defoliation can achieve staggered flowering times in these species as well. This may be a general feature of grasses, but appears to not be universal in angiosperms as early-season defoliation of photoperiod-sensitive strawberries did not affect flowering time (Guttridge 1959).

Sugar export and phloem loading mutants in maize that carry lesions in *tie-dyed1*, *tie-dyed2*, *sucrose export defective1*, *psychedelic*, and *sucrose trasporter1* have all been shown to delay flowering time (Braun *et al.* 2006; Baker and Braun 2008; Ma *et al.* 2008; Slewinski *et al.* 2009; Slewinski and Braun 2010). We also observed declines in sugar levels in *Oy1-N1989* mutants (Figure 7 and Table S11) and a delay in flowering time (Figure 1). It is tempting to consider these mutants as a demonstration that sugar signaling can work independent of chlorophyll, but these mutants also display low chlorophyll contents (Braun *et al.* 2006; Baker and Braun 2008; Ma *et al.* 2008; Slewinski *et al.* 2009; Slewinski and Braun 2010). Curiously, *sucrose export defective1* encodes tocopherol cyclase (Porfirova *et al.* 2002; Sattler *et al.* 2003). Tocopherol and chlorophyll share the phytol side chain, and the salvage pathway for phytol side chains from chlorophyll can contribute to tocopherol accumulation in both maize and Arabidopsis (Ischebeck *et al.* 2006; Schelbert *et al.* 2009; Diepenbrock *et al.* 2017). While it is attractive to try and unify these findings, chlorophyll breakdown products were only rate limiting for tocopherol synthesis in senescent tissues of Arabidopsis (Zhang *et al.* 2014). Furthermore, in *Oy1-N1989* mutants the phytol precursors should be abundant as this pool is not being consumed by chlorophyll biosynthesis. Future experiments that more carefully explore these metabolites, for instance, via other mutants that do not simultaneously affect multiple pathways, especially chlorophyll metabolism, are required.

Accumulation of sugars in the leaves due to sink disruption has been proposed to induce leaf senescence in a variety of angiosperms, including maize and Arabidopsis (Allison and Weinmann 1970; Ceppi *et al.* 1987; Pourtau *et al.* 2006; Sekhon *et al.* 2012, 2019). In maize, leaf senescence is triggered when pollination is prevented and sucrose accumulates in leaves due to the lack of the sink activity of a pollinated ear (Allison and Weinmann 1970; Ceppi *et al.* 1987). This sucrose-dependent leaf senescence is genotype-dependent, and B73 is particularly susceptible to this phenomenon. Genetic inheritance of the induced senescence phenomenon in maize inbred line B73 was proposed to be under the control of a single dominant locus (Ceppi *et al.* 1987). The leaf senescent phenotypes of the sucrose export mutants demonstrate that excessive photosynthate accumulation can cause tissue to senesce regardless of ear presence (Braun *et al.* 2006; Baker and Braun 2008; Ma *et al.* 2008; Slewinski *et al.* 2009). Induced leaf senescence by sink removal or sugar application shows some overlap of biochemical and molecular mechanisms with natural senescence in plants (summarized in Sekhon *et al*. 2012). A study of gene expression in the B73 inbred of maize identified senescence associated genes that exhibit gene expression changes during pollination-prevented leaf senescence (Sekhon *et al*. 2012). We found that leaf senescence could be prevented or delayed by the suppression of photosynthesis in the *Oy1-N1989* mutant and further modulated by *vey1* variants (Figure 9 and Table S12). We expect that future gene expression studies in *Oy1-N1989* mutant and sugar export mutants will identify genes consistently impacted by lower and higher chlorophyll levels, presumably including senescence associated genes and genes regulating carbohydrate metabolism.

## Acknowledgements

We would like to thank the National Science Foundation for NSF grant 1444503 awarded to G.S.J. and B.P.D which helped fund this research. We thank the Purdue ACRE and ICSC staff members Mr. Jim Beaty and Mr. Jason Adams for help with planting, and management of the field experiments described in this work. We also thank past Johal lab members for help with planting and data collection. We would like to thank Mr. Mike Gosney for helping with gas-exchange measurements. We gratefully acknowledge solidarity with the the operator of the compact track loader who provided a soundtrack and dry workspace by excavating and constructing a French drain on our office walls while we wrote this paper.

