## Supplemental tables 4-12 for "Interaction between induced and natural variation at *oil yellow1* delays reproductive maturity in maize"

**Allelic interactions of induced and natural variation at *oil* *yellow1* impacts reproductive maturity in maize**

### Table S4. The reproductive maturity of the wild-type (*+/+*) and mutant (*Oy1-N1989/+*) siblings in the F_1_ generation of crosses of B73 and Mo17 with *Oy1-N1989/oy1*:B73.

| Genotype | DTA | DTS | ASI |
| --- | --- | --- | --- |
| *+*/*+*:B73 | 59.6±1.4^b^ | 61.6±1.6^b^ | -2.0±0^ab^ |
| *Oy1-N1989/+*:B73 | 63.7±1.6^c^ | 65.1±1.2^c^ | -1.4±0.5^a^ |
| *+*/*+*:Mo17/B73 | 53.6±1.2^a^ | 56±1.4^a^ | -2.4±0.9^b^ |
| *Oy1-N1989/+*:Mo17/B73 | 64±1.4^c^ | 65.6±1.2^c^ | -1.6±0.9^a^ |

The data are represented as means ± SD. The values are means of 10 replicates planted in RCBD.The superscript connecting letter report between genotypes within each trait was determined using ANOVA with post-hoc analysis using Tukey’s HSD at p<0.01.

### Table S5. The trait correlations in IBM x *Oy1-N1989/+*:B73 F_1_ test cross populations.

|  | WT_CCMI | WT_CCMII | MT_CCMI | MT_CCMII | Ratio_CCMI | Ratio_CCMII | WT_DTA | WT_DTS | WT_ASI | MT_DTA | MT_DTS | MT_ASI | Ratio_DTA | Ratio_DTS | Diff_DTA | Diff_DTS | Diff_ASI |
| --- | --- | --- | --- | --- | --- | --- | --- | --- | --- | --- | --- | --- | --- | --- | --- | --- | --- |
| WT_CCMI |  | 0.3683 | 0.0691 | 0.0624 | -0.2217 | -0.0447 | -0.0899 | -0.0675 | -0.0764 | -0.0476 | -0.0551 | 0.0254 | 0.0157 | -0.0095 | -0.0084 | 0.0137 | -0.0786 |
| WT_CCMII | <.0001 |  | 0.0588 | 0.1361 | -0.0622 | -0.1564 | -0.2388 | -0.2449 | -0.0283 | -0.0992 | -0.1019 | -0.0046 | 0.0642 | 0.0583 | -0.0485 | -0.0453 | -0.0184 |
| MT_CCMI | 0.3122 | 0.3902 |  | 0.875 | 0.9475 | 0.859 | -0.0239 | -0.0464 | 0.0557 | -0.8239 | -0.8275 | -0.1224 | -0.7567 | -0.7526 | 0.7814 | 0.7745 | 0.1367 |
| MT_CCMII | 0.3614 | 0.0457 | <.0001 |  | 0.8286 | 0.9472 | -0.0171 | -0.0428 | 0.0653 | -0.7981 | -0.7977 | -0.1361 | -0.7358 | -0.726 | 0.7605 | 0.7477 | 0.1545 |
| Ratio_CCMI | 0.001 | 0.3629 | <.0001 | <.0001 |  | 0.8549 | -0.0063 | -0.0299 | 0.0618 | -0.7919 | -0.7892 | -0.1451 | -0.7384 | -0.727 | 0.7611 | 0.7471 | 0.1588 |
| Ratio_CCMII | 0.5135 | 0.0215 | <.0001 | <.0001 | <.0001 |  | 0.0476 | 0.0241 | 0.0716 | -0.774 | -0.7708 | -0.1446 | -0.7552 | -0.7423 | 0.7764 | 0.761 | 0.1659 |
| WT_DTA | 0.1882 | 0.0004 | 0.727 | 0.8026 | 0.9266 | 0.4865 |  | 0.9366 | 0.356 | 0.2554 | 0.2132 | 0.2308 | -0.4239 | -0.3973 | 0.3578 | 0.3442 | 0.0993 |
| WT_DTS | 0.3233 | 0.0003 | 0.4972 | 0.5318 | 0.6617 | 0.7247 | <.0001 |  | 0.006 | 0.2816 | 0.2498 | 0.1888 | -0.3597 | -0.4066 | 0.2942 | 0.346 | -0.1396 |
| WT_ASI | 0.2633 | 0.6792 | 0.4154 | 0.3398 | 0.3657 | 0.2948 | <.0001 | 0.9298 |  | -0.0221 | -0.0577 | 0.1551 | -0.2504 | -0.0495 | 0.2365 | 0.0595 | 0.6558 |
| MT_DTA | 0.4866 | 0.1462 | <.0001 | <.0001 | <.0001 | <.0001 | 0.0001 | <.0001 | 0.747 |  | 0.9764 | 0.2732 | 0.7666 | 0.741 | -0.8115 | -0.7805 | -0.2259 |
| MT_DTS | 0.4206 | 0.1355 | <.0001 | <.0001 | <.0001 | <.0001 | 0.0016 | 0.0002 | 0.3985 | <.0001 |  | 0.059 | 0.7713 | 0.7823 | -0.8142 | -0.822 | -0.0898 |
| MT_ASI | 0.7107 | 0.9459 | 0.0725 | 0.0457 | 0.033 | 0.0337 | 0.0006 | 0.0054 | 0.0226 | <.0001 | 0.3883 |  | 0.108 | -0.0594 | -0.1244 | 0.0539 | -0.6441 |
| Ratio_DTA | 0.819 | 0.3475 | <.0001 | <.0001 | <.0001 | <.0001 | <.0001 | <.0001 | 0.0002 | <.0001 | <.0001 | 0.1135 |  | 0.9586 | -0.9966 | -0.9588 | -0.2764 |
| Ratio_DTS | 0.89 | 0.3938 | <.0001 | <.0001 | <.0001 | <.0001 | <.0001 | <.0001 | 0.4697 | <.0001 | <.0001 | 0.3854 | <.0001 |  | -0.9558 | -0.9971 | 0.0071 |
| Diff_DTA | 0.9027 | 0.478 | <.0001 | <.0001 | <.0001 | <.0001 | <.0001 | <.0001 | 0.0005 | <.0001 | <.0001 | 0.068 | <.0001 | <.0001 |  | 0.9619 | 0.2782 |
| Diff_DTS | 0.8418 | 0.5079 | <.0001 | <.0001 | <.0001 | <.0001 | <.0001 | <.0001 | 0.3844 | <.0001 | <.0001 | 0.4308 | <.0001 | <.0001 | <.0001 |  | 0.0049 |
| Diff_ASI | 0.2501 | 0.7884 | 0.0448 | 0.0231 | 0.0196 | 0.0146 | 0.1459 | 0.0404 | <.0001 | 0.0008 | 0.1887 | <.0001 | <.0001 | 0.9178 | <.0001 | 0.9431 |  |

Upper half contains Pearson’s correlation coefficient and lower half contains correlation p-values for each pairwise trait comparison. Self-comparisons (diagonal) are left blank.

### Table S6. The trait correlations in Syn10 x *Oy1-N1989/+*:B73 F_1_ test cross populations.

|  | WT_CCMI | WT_CCMII | MT_CCMI | MT_CCMII | Ratio_CCMI | Ratio_CCMII | WT_DTA | WT_DTS | WT_ASI | MT_DTA | MT_DTS | MT_ASI | Ratio_DTA | Ratio_DTS | Diff_DTA | Diff_DTS | Diff_ASI |
| --- | --- | --- | --- | --- | --- | --- | --- | --- | --- | --- | --- | --- | --- | --- | --- | --- | --- |
| WT_CCMI |  | 0.2932 | 0.127 | 0.1175 | -0.1394 | 0.0785 | -0.1994 | -0.1951 | 0.0302 | -0.0835 | -0.0981 | 0.0942 | -0.0215 | -0.0326 | 0.0305 | 0.042 | -0.0625 |
| WT_CCMII | <.0001 |  | 0.0085 | 0.0055 | -0.0611 | -0.1443 | -0.0489 | -0.0574 | 0.0264 | 0.0289 | 0.0331 | -0.0277 | 0.045 | 0.0524 | -0.043 | -0.0506 | 0.0444 |
| MT_CCMI | 0.0448 | 0.893 |  | 0.9114 | 0.9567 | 0.8972 | -0.0466 | -0.0446 | 0.0054 | -0.8768 | -0.8657 | 0.0665 | -0.8741 | -0.8585 | 0.8789 | 0.8638 | -0.0562 |
| MT_CCMII | 0.0636 | 0.9306 | <.0001 |  | 0.8734 | 0.9853 | -0.0292 | -0.0385 | 0.0243 | -0.9002 | -0.8903 | 0.0767 | -0.9031 | -0.8851 | 0.9075 | 0.8906 | -0.0514 |
| Ratio_CCMI | 0.0276 | 0.3361 | <.0001 | <.0001 |  | 0.8716 | -0.0052 | -0.0057 | 0.0019 | -0.8483 | -0.8343 | 0.0459 | -0.8579 | -0.8396 | 0.8612 | 0.8435 | -0.0402 |
| Ratio_CCMII | 0.2163 | 0.0225 | <.0001 | <.0001 | <.0001 |  | -0.0279 | -0.0342 | 0.0181 | -0.8933 | -0.8851 | 0.085 | -0.8964 | -0.8811 | 0.9008 | 0.8865 | -0.0636 |
| WT_DTA | 0.0015 | 0.4416 | 0.4637 | 0.6454 | 0.9343 | 0.6604 |  | 0.8654 | 0.0736 | 0.1956 | 0.1982 | -0.0433 | -0.107 | -0.0845 | 0.0735 | 0.0533 | 0.0902 |
| WT_DTS | 0.0019 | 0.3664 | 0.4826 | 0.5443 | 0.9291 | 0.5904 | <.0001 |  | -0.436 | 0.1615 | 0.1889 | -0.1779 | -0.1012 | -0.1384 | 0.0714 | 0.1024 | -0.1667 |
| WT_ASI | 0.635 | 0.6779 | 0.932 | 0.7023 | 0.9765 | 0.7763 | 0.2461 | <.0001 |  | 0.0287 | -0.0212 | 0.2762 | 0.0097 | 0.124 | -0.0105 | -0.1082 | 0.4927 |
| MT_DTA | 0.1883 | 0.6493 | <.0001 | <.0001 | <.0001 | <.0001 | 0.0019 | 0.0106 | 0.6514 |  | 0.9841 | -0.0574 | 0.9539 | 0.9395 | -0.9637 | -0.9495 | 0.0733 |
| MT_DTS | 0.122 | 0.6028 | <.0001 | <.0001 | <.0001 | <.0001 | 0.0016 | 0.0027 | 0.7391 | <.0001 |  | -0.2338 | 0.9367 | 0.9462 | -0.9468 | -0.9575 | 0.196 |
| MT_ASI | 0.1375 | 0.6633 | 0.2949 | 0.2267 | 0.4696 | 0.1801 | 0.4957 | 0.0048 | <.0001 | 0.3659 | 0.0002 |  | -0.0432 | -0.1756 | 0.0466 | 0.1846 | -0.7003 |
| Ratio_DTA | 0.7351 | 0.4788 | <.0001 | <.0001 | <.0001 | <.0001 | 0.0913 | 0.1105 | 0.8782 | <.0001 | <.0001 | 0.497 |  | 0.9784 | -0.9992 | -0.9786 | 0.0463 |
| Ratio_DTS | 0.608 | 0.4095 | <.0001 | <.0001 | <.0001 | <.0001 | 0.1831 | 0.0287 | 0.0502 | <.0001 | <.0001 | 0.0054 | <.0001 |  | -0.9784 | -0.9991 | 0.251 |
| Diff_DTA | 0.631 | 0.4981 | <.0001 | <.0001 | <.0001 | <.0001 | 0.247 | 0.2609 | 0.8691 | <.0001 | <.0001 | 0.4632 | <.0001 | <.0001 |  | 0.9801 | -0.05 |
| Diff_DTS | 0.5081 | 0.4258 | <.0001 | <.0001 | <.0001 | <.0001 | 0.4011 | 0.1062 | 0.0878 | <.0001 | <.0001 | 0.0034 | <.0001 | <.0001 | <.0001 |  | -0.2475 |
| Diff_ASI | 0.3249 | 0.485 | 0.3764 | 0.4182 | 0.5269 | 0.3166 | 0.1548 | 0.0083 | <.0001 | 0.2481 | 0.0019 | <.0001 | 0.4661 | <.0001 | 0.4314 | <.0001 |  |

Upper half contains Pearson’s correlation coefficient and lower half contains correlation p-values for each pairwise trait comparison. Self-comparisons (diagonal) are left blank.

### Table S7. The summary of the QTL detected for reproductive maturity in IBM x *Oy1-N1989/+*:B73 F_1_ test cross populations.

|  |  |  |  |  |  | LOD-2 Interval^e^ | |  | Mean±SE^g^ | |  |
| --- | --- | --- | --- | --- | --- | --- | --- | --- | --- | --- | --- |
| Trait Identifier | Chr^b^ | LOD^c^ | Position^d^ | Left Marker | Right Marker | Left | Right | PVE^f^ | B73 | Mo17 | Greater Allele |
| MT_DTA | 10 | 29.7 | 144 | phi059 | isu85b | 124 | 146.8 | 46.92 | 69.58±0.27 | 75.31±0.33 | Mo17 |
| MT_DTS | 10 | 33.8 | 127 | umc2034 | AI795367 | 124 | 131 | 39.50 | 69.97±0.29 | 74.70±0.39 | Mo17 |
| Ratio_DTA | 10 | 32.9 | 139 | AI795367 | AY109994 | 128 | 146 | 47.04 | 1.11±0.01 | 1.21±0.01 | Mo17 |
| Ratio_DTS | 10 | 32.1 | 139 | AI795367 | AY109994 | 125 | 146 | 46.60 | 1.08±0.01 | 1.17±0.01 | Mo17 |
| Diff_DTA | 10 | 34.6 | 139 | AI795367 | AY109994 | 127 | 146 | 48.39 | -7.01±0.28 | -12.88±0.35 | B73 |
| Diff_DTS | 10 | 34.0 | 129 | umc2034 | AI795367 | 124 | 146 | 45.76 | -5.87±0.27 | -11.36±0.37 | B73 |

^a^Number of QTL detected for a given trait

^b^Chromosome location of a QTL

^c^LOD score at the peak of a given QTL

^d^Peak position and ^e^LOD-2 interval of the QTL in terms of genetic position in centiMorgan(cM)

^f^PVE is percent of variance explained by the QTL at this position as estimated by regression and reported as an R^2^ value*100

^g^Mean and standard error of the trait with B73 and Mo17 genotype at the peak marker of the detected QTL

### Table S8. The summary of the QTL detected for reproductive maturity in Syn10 DH x *Oy1-N1989/+*:B73 F_1_ test cross populations.

|  |  |  |  |  |  |  | LOD-2 Interval^e^ | |  | Mean±SE^g^ | |  |
| --- | --- | --- | --- | --- | --- | --- | --- | --- | --- | --- | --- | --- |
| Trait Identifier | QTL^a^ | Chr^b^ | LOD^c^ | Position^d^ | Left Marker | Right Marker | Left | Right | PVE^f^ | B73 | Mo17 | Greater Allele |
| MT_DTA | 2 | 2 | 4.4 | 619.7 | chr02.970 | chr02.1038 | 613.6 | 757 | 7.70 | 62.27±0.33 | 64.82±0.45 | Mo17 |
|  |  | 10 | 65.7 | 192 | chr10.94.5 | chr10.95.5 | 188.7 | 193 | 66.09 | 60.8±0.18 | 68.73±0.29 | Mo17 |
| MT_DTS | 2 | 2 | 4.2 | 619.7 | chr02.970 | chr02.1038 | 613.6 | 755.8 | 7.36 | 63.52±0.34 | 66.07±0.46 | Mo17 |
|  |  | 10 | 64.4 | 192 | chr10.94.5 | chr10.95.5 | 188.7 | 193 | 66.10 | 61.95±0.19 | 70.1±0.29 | Mo17 |
| Ratio_DTA | 1 | 10 | 71.4 | 192 | chr10.94.5 | chr10.95.5 | 190.5 | 193 | 67.56 | 1.08±0.003 | 1.22±0.004 | Mo17 |
| Ratio_DTS | 1 | 10 | 66.7 | 192 | chr10.94.5 | chr10.95.5 | 190.5 | 193 | 65.95 | 1.07±0.003 | 1.21±0.005 | Mo17 |
| Diff_DTA | 1 | 10 | 73.3 | 192 | chr10.94.5 | chr10.95.5 | 190.5 | 193 | 68.12 | -4.75±0.18 | -12.69±0.27 | B73 |
| Diff_DTS | 1 | 10 | 69.1 | 192 | chr10.94.5 | chr10.95.5 | 190.5 | 193 | 66.88 | -4.48±0.19 | -12.56±0.29 | B73 |

^a^Number of QTL detected for a given trait

^b^Chromosome location of a QTL

^c^LOD score at the peak of a given QTL

^d^Peak position and ^e^LOD-2 interval of the QTL in terms of genetic position in centiMorgan(cM)

^f^PVE is percent of variance explained by the QTL at this position as estimated by regression and reported as an R^2^ value*100

^g^Mean and standard error of the trait with B73 and Mo17 genotype at the peak marker of the detected QTL

### Table S9. The linear regression of various flowering traits in IBM X *Oy1*-*N1989*/*+*:B73 F_1_ progenies on to the normalized OY1 expression data (RPKM) derived from shoot apices of 14 days old corresponding IBM-RILs (n=74) seedlings.

| Trait/Marker | R^2^ (%) | P-value |
| --- | --- | --- |
| isu085b | 19.2 | <0.0001 |
| MT_DTA | 20.7 | <0.0001 |
| WT_DTA | 0.19 | 0.96 |
| MT_DTS | 18.2 | <0.0002 |
| WT_DTS | 0.02 | 0.90 |
| Ratio_DTA | 17.5 | <0.0002 |
| Ratio_DTS | 14.7 | <0.0007 |
| Diff_DTA | 18.8 | <0.0001 |
| Diff_DTS | 15.4 | <0.0006 |
| MT_ASI | 1.2 | 0.34 |
| WT_ASI | 0.1 | 0.81 |
| Diff_ASI | 1.4 | 0.31 |

### Table S10. The allele specific expression analysis of the OY1 transcript in the leaf tissue from the top collared leaf at V3 developmental stage derived from the F_1_ hybrids between B73-like NILs and *Oy1*-*N1989*/*+*:B73.

| Pedigree | *vey1*-status | Ref_allele^1^ | Alt_allele^2^ | Ref_depth^3^ | Alt_depth^4^ | Allele_Ratio^5^ |
| --- | --- | --- | --- | --- | --- | --- |
| *Oy1*-*N1989*/*+*:b135/B73 | *vey1^B73^* | C | T | 1130.7±55.1 | 1307.7±92.8 | 1.16±0.03^a^ |
| *Oy1*-*N1989*/*+*:b185/B73 | *vey1^B73^* | C | T | 909.7±174.9 | 1063.3±220.9 | 1.17±0.03^a^ |
| *Oy1*-*N1989*/*+*:b094/B73 | *vey1^Mo17^* | C | T | 702±1.7 | 978.3±55.3 | 1.39±0.08^b^ |
| *Oy1*-*N1989*/*+*:b189/B73 | *vey1^Mo17^* | C | T | 622.3±147.4 | 884.7±190.1 | 1.43±0.12^b^ |

The data is provided as mean and standard deviations. The superscript connecting letter report for last column was determined using ANOVA with post-hoc analysis using Tukey’s HSD at p<0.05.

^1,2^SNP genotype at the causative mutation of *Oy1*-*N1989* allele (L176F) representing the wild-type/reference (Ref_allele) and mutant/alternate (Alt_allele) allele.

^3,4^Sequence depth of the high quality reads at the measured SNP for reference and alternate allele.

^5^Ratio of the high quality sequence reads at the measured SNP (Alternate/Reference).

### Table S11. The profiling of soluble and insoluble carbohydrates in the top fully-expanded leaf at V3 developmental stage in field-grown maize seedlings.

| Pedigree | *vey1*-status | Fructose | Glucose | Sucrose | Starch |
| --- | --- | --- | --- | --- | --- |
| *Oy1*-*N1989*/*+*:b135/B73 | *vey1^B73^* | 0.11±0^a^ | 0.77±0.09^a^ | 4.65±0.65^a^ | 5.09±0.51^a^ |
| *Oy1*-*N1989*/*+*:b185/B73 | *vey1^B73^* | 0.14±0.03^a^ | 0.76±0.06^a^ | 4.13±0.68^a^ | 4.42±0.50^a^ |
| *Oy1*-*N1989*/*+*:b094/B73 | *vey1^Mo17^* | 0.06±0.01^b^ | 0.44±0.08^b^ | 1.31±0.33^b^ | 0.66±0.25^b^ |
| *Oy1*-*N1989*/*+*:b189/B73 | *vey1^Mo17^* | 0.06±0.01^b^ | 0.52±0.01^b^ | 1.69±0.14^b^ | 0.81±0.09^b^ |

The data are presented as means and standard deviation derived from three replications planted in RCBD. All measurements are expressed as mg/g of fresh weight. Each biological replication consists of tissue pooled from 3-4 independent plants. The connecting letter report indicates statistical significance determined using ANOVA followed by mean comparisons using Tukey’s HSD at p<0.05.

### Table S12. The visual quantification of leaf senescence of primary ear leaf induced by pollination prevention in B73-like NILs × *Oy1-N1989/+*:B73 F_1_ progenies.

| Genotype | *vey1*-status | 30DAA | 35DAA | 40DAA | 45DAA |
| --- | --- | --- | --- | --- | --- |
| *+*/*+* | *vey1^B73^* | 7.4±1.7^a^ | 9±1.2^a^ | 9.9±0.3^a^ | 10±0^a^ |
| *Oy1*-*N1989*/*+* | *vey1^B73^* | 3.3±2.0^b^ | 6.4±1.5^b^ | 8.6±1.3^b^ | 10±0^a^ |
| *+*/*+* | *vey1^Mo17^* | 7.9±1.4^a^ | 9.7±0.6^a^ | 10±0^a^ | 10±0^a^ |
| *Oy1*-*N1989*/*+* | *vey1^Mo17^* | 0.1±0.3^c^ | 0.4±0.7^c^ | 2.6±1.2^c^ | 6.4±0.8^b^ |

The data are presented as means ± standard deviation derived from three replications planted in RCBD of eight (*vey1^Mo17^* allele) and fifteen (*vey1^B73^* allele) independent NIL F_1_ progenies. The visual assessments of leaf greenness were done on a scale of 0-10, with 0 being 100% green leaf and 10 being totally dry leaf. The visual scoring was done at 5-day interval from 30-45 days after anthesis (DAA) on per plot basis using plants with non-pollinated primary and secondary ears. The connecting letter report indicates statistical significance determined using ANOVA followed by mean comparisons using Tukey’s HSD at p<0.05.
